# A large-scale and long-term experiment to identify effectiveness of ecosystem restoration

**DOI:** 10.1101/2024.04.02.587693

**Authors:** Merja Elo, Santtu Kareksela, Otso Ovaskainen, Nerea Abrego, Jenni Niku, Sara Taskinen, Kaisu Aapala, Janne S. Kotiaho

## Abstract

Ecosystem restoration will increase following the ambitious international targets, which calls for a rigorous evaluation of restoration effectiveness. Studies addressing restoration effectiveness across ecosystems have thus far shown varying and unpredictable patterns. A rigorous assessment of the factors influencing restoration effectiveness is best done with large-scale and long-term experimental data. Here, we present results from a well replicated long-term before-after control-impact experiment on restoration of forestry-drained boreal peatland ecosystems. Our data comprise 151 sites, representing six ecosystem types. Vegetation sampling has been conducted to the species level before restoration and two, five and ten years after restoration. We show that, on average, restoration stops and reverses the trend of further degradation. The variation in restoration outcomes largely arises from ecosystem types: restoration of nutrient-poor ecosystems has higher probability of failure. Our experiment provides clear evidence that restoration can be effective in halting the biodiversity loss of degraded ecosystems, although ecosystem attributes can affect the restoration outcome. These findings underlie the need for evidence-based prioritization of restoration efforts across ecosystems.

## INTRODUCTION

Ecosystem restoration is likely to become mainstream in the near future, even if the Kunming-Montréal Global Biodiversity Framework target ^1^ to effectively restore at least 30% of degraded ecosystems by 2030 is so ambitious that it is unlikely to be met. As 20-40% of the global land area is degraded, even smaller percentage of the land to be restored would add to the 1 billion hectares currently committed to restoration ^2^. In general, restoration succeeds in increasing biodiversity and ecosystem functions in degraded ecosystems ^3–5^. Yet, restored ecosystems rarely fully recover ^4–6^, and the outcomes are often unpredictable ^3,5,7^. To make restoration more effective, we need to improve the predictability of restoration by identifying the causal factors behind the variation in the restoration outcomes ^7,8^.

Identification of the sources for the variation is best done with properly controlled and replicated experimental data ^9^. Unfortunately, such data are relatively scarce in environmental biology. Only 23% of the biological intervention studies use randomised designs or controlled observational designs with before impact sampling (i.e., before-after control-impact -design), despite the fact, that such designs are known to provide less biased estimates than simpler designs ^9^. In restoration studies, the amount of experimental studies is even lower. A recent meta-analysis of restoration effects ^5^ included 89 studies of which only ten had before-after control-impact -design, and only half had both unrestored and reference controls which are needed to estimate and verify the effect of restoration reliably. Thus, the repeated calls for properly controlled and long-term restoration experiments, replicated over large spatial scales ^8,10^ remain to be answered.

Here, we contribute to filling this gap by reporting results from a well replicated long-term before- after control-impact experiment on restoration of forestry-drained boreal peatland ecosystems (Fig. 1). These ecosystems rely on high water table level and are thus important carbon sinks and storages^11,12^, provide numerous other ecosystem services ^13^, and host unique biodiversity ^14^ in their pristine stage. Boreal peatland ecosystems have been widely drained for agriculture, peat extraction and especially for forestry, the latter affecting approximately 30% of Europe’s peatland ecosystem area ^15^. Promisingly, the forestry-drained ecosystems may be more easily restored in comparison to the other uses because the vegetation is not removed, and there are likely to be source populations nearby for species that have become locally absent ^16,17^. For instance, in Finland alone there are up to one million hectares of mire forests and fens where the drainage has not been successful in increasing the tree growth to a profitable level ^18^, mainly due to lack or imbalance of nutrients in these sites. Hence, restoring forestry-drained ecosystems could result in long-term biodiversity ^14^ and climate benefits ^19^.

**Fig. 1.**
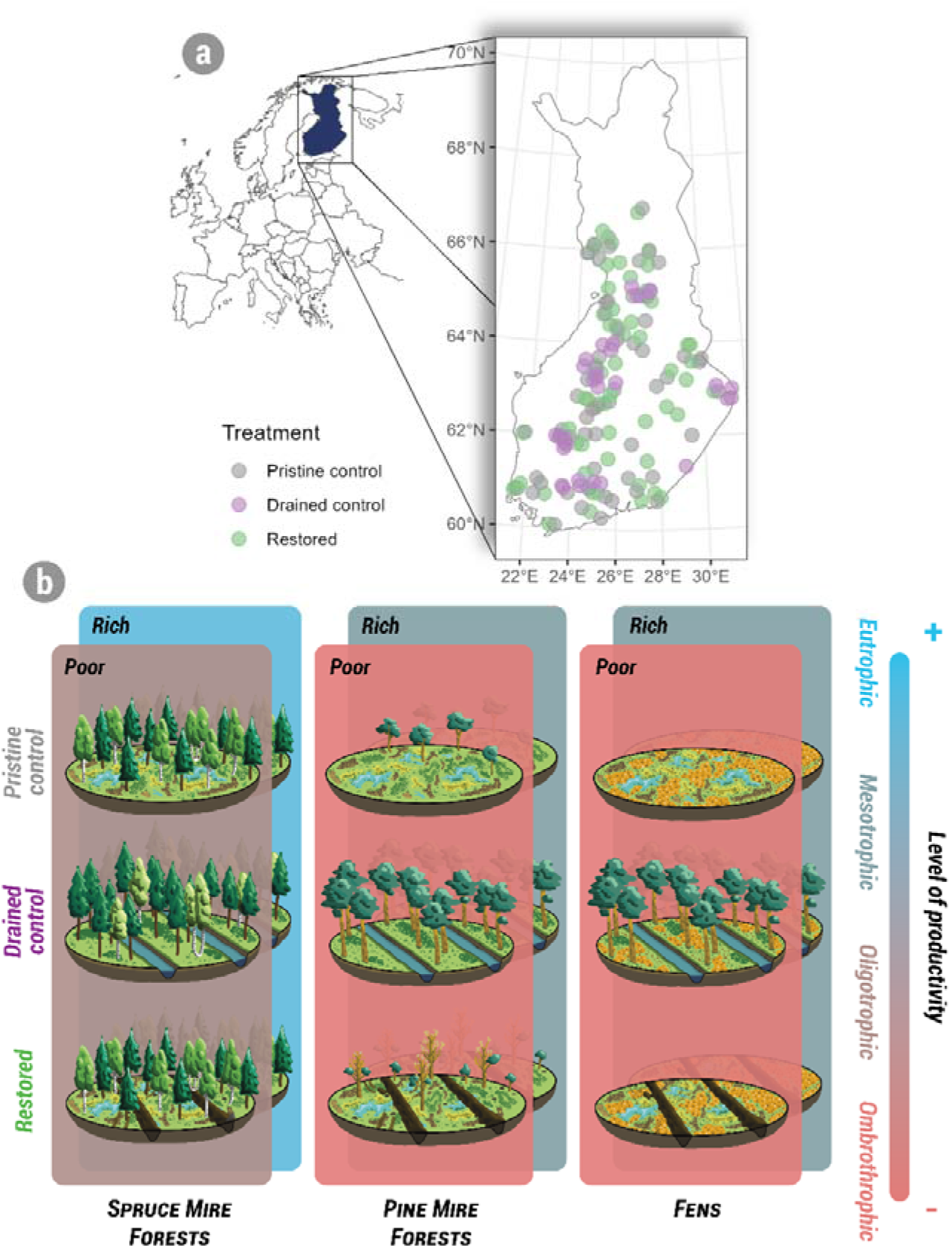
Map of the 151 study sites in Finland: sites that had been drained for forestry during 1960s and 1970s and were restored during the project (‘restored’; green symbols), relatively pristine sites with no drainage (‘pristine control’; grey symbols), and sites that had been drained and were not restored (‘drained control’; purple symbols) (a). The sites represent three main ecosystem types: spruce mire forests, pine mire forests, and fens, which all rely on high water table level. They are further divided to two types (poor/rich) according to the productivity level (as shown by the two overlapping panels; note that the productivity levels ‘poor’ and ‘rich’ are not comparable between spruce mire forests, pine mire forests and fens) (b). Hence, the six ecosystem types form a rough gradient in their productivity level and differ in their tree cover and species composition. Each site includes ten permanent sampling plots from which vascular plant and moss species were monitored prior to restoration and two, five and ten years after restoration in restored, drained control and pristine control sites (Fig. S1)

Indeed, restoration of forestry-drained ecosystems often succeeds in rising the water table to pristine levels ^20,21^. This often leads to recovery of the ecosystem functions, such as carbon sequestration ^22–24^, but only a partial recovery of community composition ^17,21,22,25–27^. However, similarly to the restoration studies in general, the previous studies are based on few study sites ^21,26^, early years after restoration ^27^ or space-for-time substitutions ^17,22,25^. Moreover, the studies have often focused on only one of the many peatland ecosystem types which may respond differently to restoration ^23^, due to their inherent differences in nutrient levels ^28^.

We show that without restoration, the forestry-drained boreal peatland ecosystems tended to continue to degrade during the monitored ten years. On average, restoration of these ecosystems was successful in stopping and reversing the trend of degradation of the vegetation communities and reversing the successional process. Our results reinforce the previous findings on the restoration effects in general ^4–6^ as well as for the boreal forestry-drained peatland ecosystems ^17,21,22,25–27^: restoration is clearly effective in changing the community composition towards, but not all the way to, the reference pristine community composition. However, our modelling revealed important variation in the responses of the vegetation communities between different ecosystem types. That is, restoration led to different outcomes depending on the ecosystem type. These results have profound implications for restoration planning when considering the probability of restoration success and when setting restoration priorities. The data-based evidence provided by our experiment can significantly contribute to improving the effectiveness of restoration.

## RESULTS AND DISCUSSION

### Overall response to restoration

In general, restoration was successful in reversing the effects of drainage: the average vegetation community composition in restored sites was more similar to pristine controls and less similar to drained controls ten years after restoration, than before restoration (Fig. 2a). This resulted from a positive response to restoration of species which had decreased after drainage, and conversely a negative response of species which had increased after drainage (Fig. 3). For instance, several peat- forming *Sphagnum* mosses responded positively whereas common forest mosses (*Hylocomnium splendens, Pleurozium schreberi*) which tend to colonize or increase in drained peatland ecosystems^16^, responded negatively (Fig. 4), as found also in previous studies ^17,25^. However, many of the responses were mild (Fig. 4), and ten years after restoration the species abundances did not equal those in the reference pristine controls (Fig. S6 in Supporting Information 1). Also, several species that had increased or decreased after drainage did not show detectable responses to restoration (Fig. 3). This resulted into a partial recovery of the plant communities (Fig. 2a). This suggest that our monitoring period of ten years, although being longer than in many restoration studies ^5,6^, is not enough for a full recovery of the plant communities. This is in line with previous meta-analyses suggesting that full recovery, if achieved at all, takes decades rather than years ^4,6^. While communities in restored sites will with time converge towards the pristine, most likely they will not fully resemble the pristine communities. One reason for this is that community succession is always affected by drift. Communities will develop somewhat differently due to stochasticity ^29,30^, even if identical at the initial stage. These drift effects are expected to be larger when the abundances of the species are small ^31^. Given that the abundances of many peatland-species have severely declined due to drainage, drift is likely to have a strong effect on how community develops in the restored sites.

**Fig. 2.**
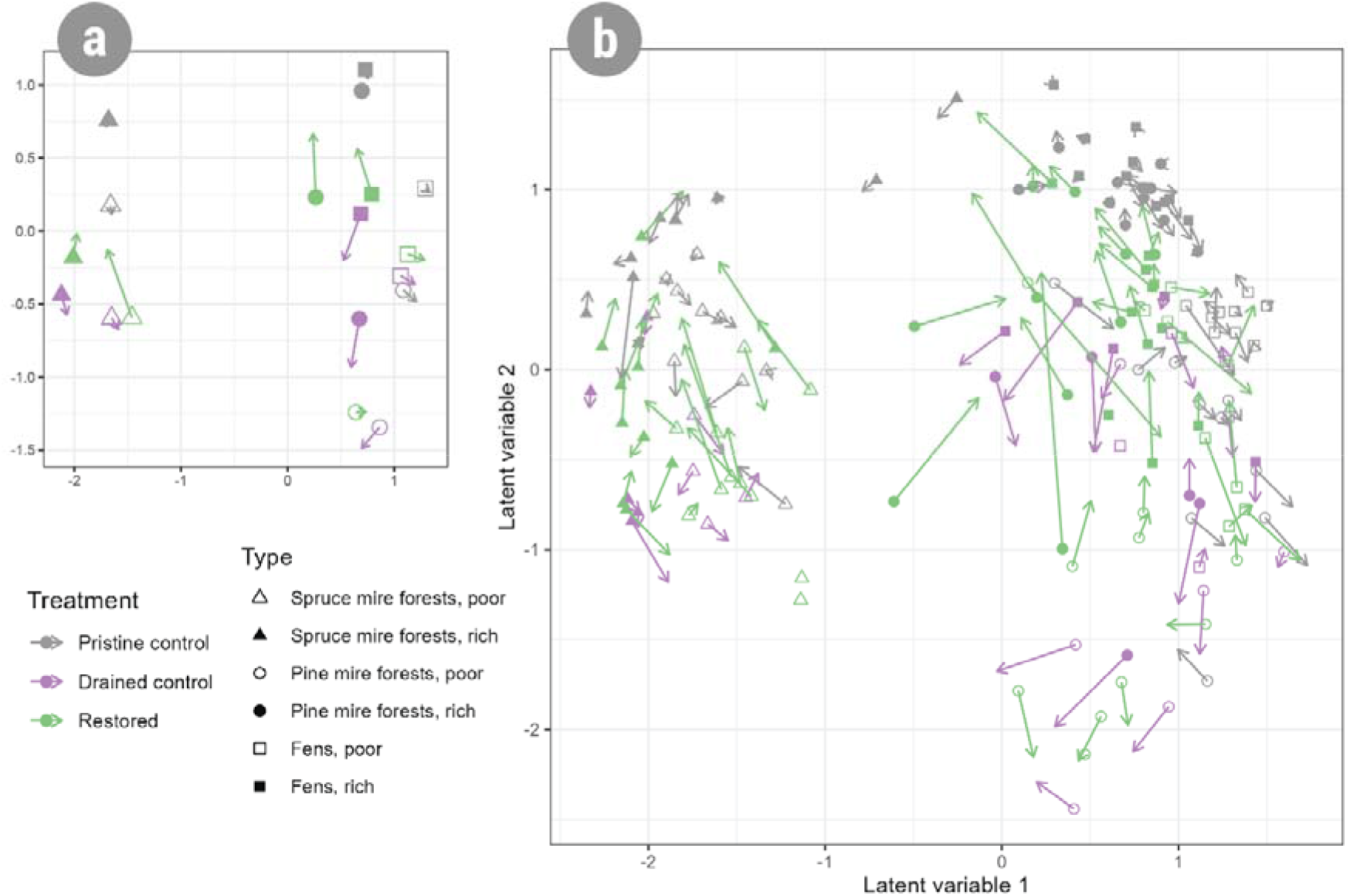
Vegetation community compositions in restored sites (green) were more similar to pristine controls (black) and less similar to drained controls (purple) 10 years after restoration (arrow head) than how they were before restoration (symbol) on average (a) but the variation between individual sites and ecosystem types was large (b). Panel **(a)** shows the average values of all sites for each treatment (restored/pristine control/drained control) per ecosystem type (poor spruce mire forest, rich spruce mire forest, poor pine mire forest, rich pine mire forest, poor fen, rich fen) from a model-based ordination before restoration (symbol) and 10 years after restoration (the head of the arrow). Panel **(b)** shows the corresponding values for each of the 151 sites.

**Fig. 3.**
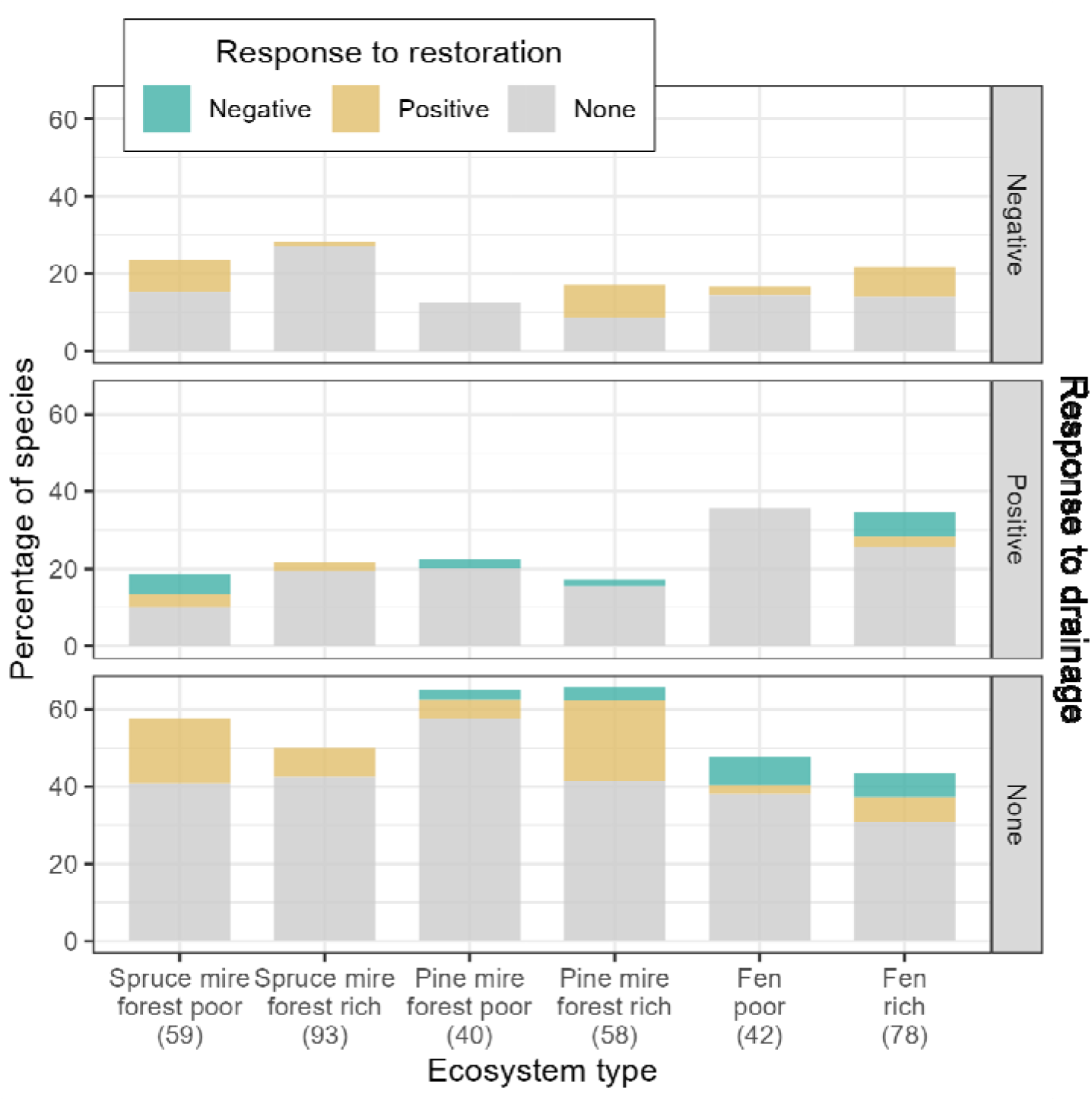
Species response to drainage and restoration. The bars show the percentage of species according to their response to drainage (negative/positive/none) and response to restoration (negative=green/positive=yellow/none=grey) in each ecosystem type (number of analysed species is shown in parentheses). The response to drainage is negative if species predicted abundance (occupancy x cover given presence, as predicted from the joint species distribution modelling) is smaller in both restored sites and drained controls than in pristine controls before restoration with high statistical support (>95% posterior probability), and positive if larger. The response to restoration is the change in abundance before restoration to ten years after restoration in restored sites in relation to drained controls. Hence, response to restoration is positive (in the figure with high statistical support >95% posterior probability) either if abundance in restored sites increases and abundance in drained sites increases less, is stable or decreases, or if abundance in drained sites decreases and abundance in restored sites is stable or decreases less. Similarly, response to restoration is negative either if abundance in restored sites decreases and abundance in drained sites decreases less, is stable or increases, or if abundance in drained sites increases and abundance in restored sites is stable or increases less.

**Fig. 4.**
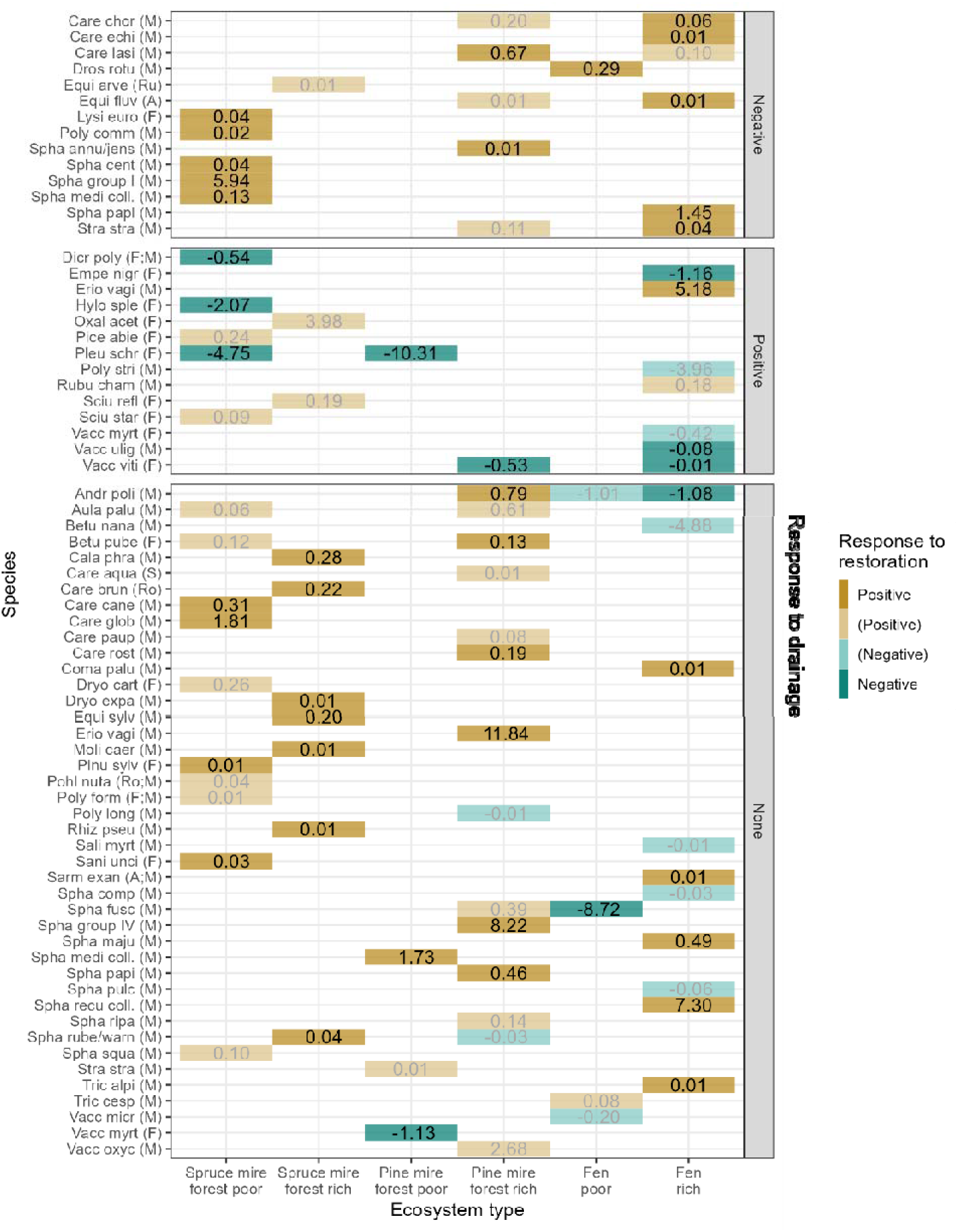
Species-specific responses to restoration. For each species responding to restoration with high statistical support (95% posterior probability) the posterior median of the response is shown, separately according to their response to drainage in the given ecosystem type. Positive responses are shown in yellow and negative in purple. The lighter colour and grey font are for species that differ in the initial abundance in drained and restored sites. For response median values between 0.01 and 0, value 0.01 is used, and for response median values between 0 and -0.01, -0.01 is used. Species full names can be found from Table S2. Species primary habitat type according to is shown in parentheses [F = forests, M = mires (including mire forests and fens), A = aquatic habitats, S = shores, Ro = rock outcrops and boulder fields, Ru = rural biotopes and cultural habitats].

Some species which did not respond or even increased after drainage responded positively to restoration (Fig. 3), possibly because they benefit from bare peat surface, increased light availability and/or the nutrient pulse due to restoration actions ^17^. For example, a sedge *Eriophorum vaginatum* increased strongly (Fig. 4), in concert with the increased nutrient release ^32^, as observed also previously ^17,23^. Likewise, birch (*Betula pubescens*) increased due to restoration (Fig. 4). In spruce mire forests, birch is a natural part of the ecosystem and tend to increase as spruce die and light penetrates to the ground ^25^ but in pine mire forests it may counteract restoration as it shades understorey and evaporates water. From mosses, especially *Sphagnum* species responded positively, most likely because they benefit from wetter conditions, with the exception of *S. fuscum* occupying drier microhabitats and consequently decreasing after restoration. Many of the positive responses were for species that differed already in their initial abundances in restored and drained sites (Fig. 4). Hence, their changes may indicate site-specific differences and may not be generalized.

Although many of the species increased above the levels of pristine controls (Fig. S6), the responses were mild and restoration effects in general tend to attenuate with time ^4,6^. Thus, pronounced community divergences are not expected in the future. Ten years after restoration, the community compositions of restored sites represent the continuum from drained, dryer heath forest type communities towards communities in pristine ecosystems (Fig. 2; Fig. 4), without for instance non-native species. Thus, by contrast to what has been suggested for restored fens in Central Europe ^33^, restoration of boreal forestry-drained peatland ecosystems do not appear to lead to self-sustaining novel ecosystems (*sensu* Hobbs *et al.* ^34^) but rather to a transforming stage between pristine and degraded ecosystems.

### Variation in the responses to restoration

In all ecosystem types, there was considerable variation in response to restoration (Fig. 2b; Fig. 5). However, restoration was clearly more effective in certain ecosystem types than in others. Approximately one third of the species responded (either positively or negatively) in poor spruce mire forests, rich pine mire forests and rich fens (34%, 34%, 29% of the species showed statistically supported responses; respectively). These ecosystem types represent the medium level of nutrient availability in the continuum of boreal peatland ecosystems (Fig. 1). By contrast, only 10-12% of the species showed statistically supported responses in poor pine mire forests, poor fens and rich spruce mires (Fig. 3) which corresponds to only four species in poor pine mire forests and five in poor fens (Fig. 4). Hence, the effect of restoration was greatest at the middle of the productivity gradient (Fig. 5). It has previously been suggested that passive restoration is most likely to succeed in medium levels of productivity because resource depletion restricts species responses at low productivity sites and competition among species increases at high productivity sites ^35^. In a short-term study where two forestry-drained ecosystems were compared two years after restoration, the nutrient-rich site recovered faster than the nutrient-poor site ^23^. This is in line with the effect of drainage: relatively wet and nutrient-rich types change faster after drainage than drier and nutrient-poor types ^16^. Hence, our results support the idea that productivity level is an important factor explaining the restoration outcome. Thus, future studies on peatland restoration should consider productivity level as a key factor explaining restoration outcome.

**Fig. 5.**
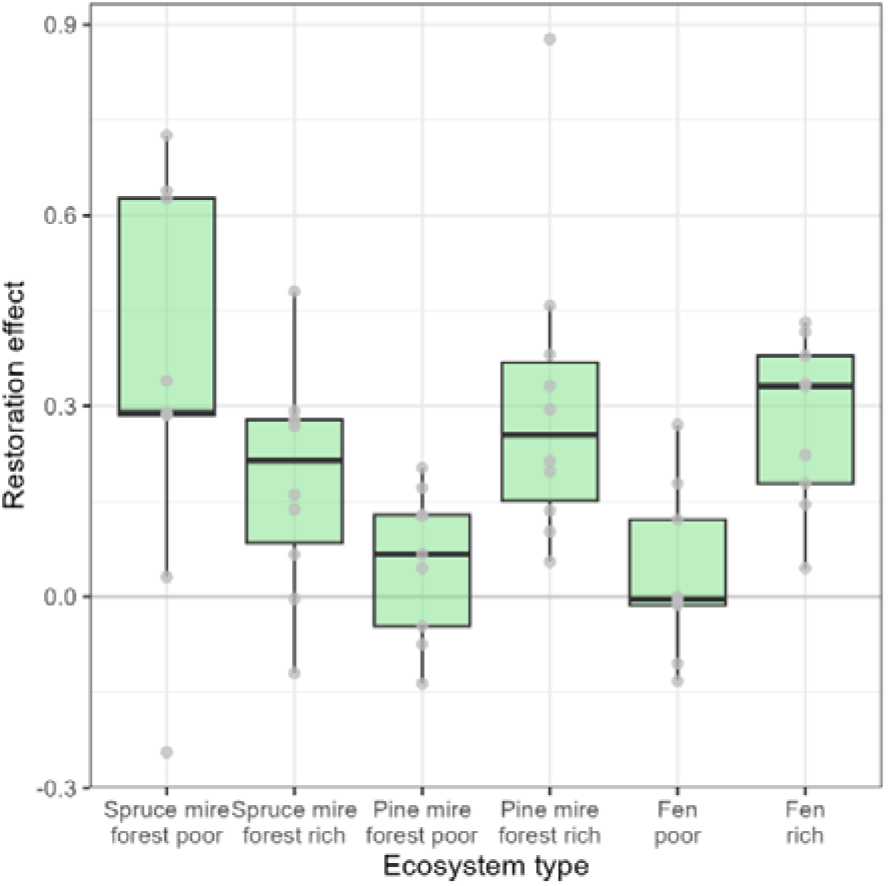
Restoration effect was variable among ecosystem types. . The restoration effect (based on the latent variables from the model-based ordination in Fig. 2) for each restored site is positive if species composition in a site changed more towards pristine sites than drained sites from before restoration to ten years after restoration. Conversely, the restoration effect is negative if species composition changed more towards drained rather than pristine sites. The thick line corresponds median, and lower and upper hinges to the 25^th^ and 75^th^ percentiles. The upper/lower whisker extends to the largest/smallest value that is at maximum 1.5 × distance between the 25^th^ and 75^th^ percentiles. Data are shown as grey points.

The negligible mean response to restoration of poor pine mire forests and fens (Fig. 2a, 3, 4) resulted partly due to opposing restoration outcomes among sites within these ecosystems (Fig. 5). For poor pine mires, some restored sites recovered well but the restored sites which had very similar communities to the drained sites before restoration, did not recover but rather followed the trajectory of the drained controls after restoration (Fig. 2b). The same but to a lesser extent applied to poor fens. Hence, the outcome of restoration may also depend on the pre-restoration community composition, which, if important but unaccounted for, can lead to unpredictable variation. These so-called priority effects are known to be important determinants of the community succession pathways ^36^ and to interact with productivity levels ^37^ In peatland ecosystems, priority effects may seem to be particularly influential for poor pine mire forests and fens. When species of drier conditions are abundant enough before restoration, they may prevent the re-colonization of the original species. In the same line, drainage can cause irreversible changes in peat hydraulic conditions, such as increase in surface compaction and peat density ^38,39^ which may prevent the pristine-like hydrological conditions in a site to recover ^40^. Furthermore, it is possible that the variation in restoration outcome was affected by the technical success of restoration. As the nutrient-poor sites are usually situated on raised bogs, receiving their water and nutrients only from rain, filling in the ditches does not necessarily result in increased water table level, in particular if the drainage has caused peat subsidence near ditches.

Regardless of the specific reason(s), increase in similarity with the pristine sites was variable and the amount of change in general small, suggesting that the restoration success in nutrient-poor pine mires and fens is uncertain. Although their restoration could be easily justified from the economic point of view, as drainage of these ecosystem types has often failed to yield economically valuable tree growth ^18^, it has lower probability to yield benefits for biodiversity when compared to nutrient- richer pine mires and fens.

### The continuing effect of drainage for forestry

The effect of restoration should be interpreted not only by the increasing similarity of the restored communities towards the pristine controls but also by their increasing dissimilarity from the drained controls ^5^. Thus, the effect of restoration is either larger or smaller than estimated from the state of the ecosystem at the time of restoration, if the unrestored sites continue to degrade or start to passively recover. Continued degradation was the case in our experiment: on average, drained sites increasingly differed from the pristine sites during the study period (Fig. 2a, Fig. S7). This resulted from the continued increase of forest mosses in poor spruce mire forests, and forest dwarf shrubs in poor pine mire forests and rich fens (Fig. S8). Altogether these changes resulted in drier, more heath forest like conditions in drained sites, highlighting that drained sites do not necessarily recover passively without restoration ^41^. Detecting the continuous degradation with our experiment is somewhat remarkable because the sites had been drained as long as 50-60 years ago, and only in three sites the ditches have been re-opened since. For rich spruce and pine mire forests, species responses behind the increasing dissimilarity from the pristine controls could not be dissected, possibly because large variation in species composition and site-specific responses (Fig. S7). By contrast, species-specific changes indicate some passive recovery.

While our experimental set-up is a large-scale and long-term well replicated before-after control-impact design, we acknowledge it also has features that affect the generalization of the results. First, it was not possible to randomize the restoration and control treatments among the drained sites. This constraint shows as a difference in initial abundances in restored treatment and drained control sites for many species (Fig. 4), which adds uncertainty to our estimates of species responses. Second, restored sites were often located in protected areas near pristine peatlands, meaning that their connectedness and pre-restoration naturalness may on average be higher compared to many drained peatlands outside protected areas and or not yet chosen for restoration. In addition, the watersheds where the study sites (both pristine and drained controls as well as restored sites) were located may be less drained in total, helping to secure the effect of the raised water table by restoration. Third, the vegetation sampling plots were situated at least 10 meters away from the ditch line. Both the effect of the drainage and the effect of the restoration are likely to be smaller and vegetation community develops more directly towards pristine communities compared to vegetation closer to or even in the ditch ^17^. In the ditch the vegetation may change fast, but not necessarily towards exactly the pristine reference community composition (often a wetter peatland type in the restored ditch) ^17^. This kind of variation in restoration outcome can however also increase the resilience of the restored peatland ecosystems through habitat heterogeneity ^17^.

Finally, we highlight that our results have also implications to understand the future changes on peatland ecosystems due to climate change. Climate change is expected to magnify the drying of peatlands because of increased temperatures and especially the more frequent and prolonged droughts ^42–44^. Hence, the effects of climate change are likely to be similar to, and possibly synergistic with, drainage for forestry, being stronger in drained sites. Even if restoration is not able to bring back original community composition, it may help in preventing further ecosystem degradation. Restoration may increase ecosystem resilience to climate change as well as mitigate climate change by preventing even further oxidation of the deeper peat layers ^45^.

### Conclusions

Restoration of forestry-drained boreal peatland ecosystems is on average effective in stopping and reversing further degradation. Our large-scale and long-term monitoring effectively revealed variation in restoration outcomes, and suggested that ecosystem characteristics, including productivity level, can significantly explain some of this variation. These results have implications for peatland restoration planning and prioritization, as well as to predict the future effects of climate change on these ecosystems. Overall, our study emphasizes the importance of large-scale and long-term restoration monitoring to guide our actions from single site planning to strategical decisions in the era of ecosystem restoration.

## METHODS

### Study system and experimental design

The data include 151 sites, located in the southern, central, and northern boreal climatic- phytogeographical zones in Finland (Fig. 1a). The sites represent six main ecosystem types: (i) poor and (ii) rich spruce mire forests, (iii) poor and (iv) rich pine mire forests, and (v) poor and (vi) rich fens. All the ecosystems are characterized by high water table level and the presence of peat-forming plant species, yet they form a rough gradient in their productivity level (note that ‘poor’ and ‘rich’ are comparable within but not between spruce mire forests/pine mire forests/fens) and differ in their tree cover and species composition (Fig. 1b).

Spruce mire forests (Fig. S2a,b; roughly corresponding EUNIS habitat type T3K ^46^) are characterised by high tree cover, dominantly spruce (*Picea abies*) in poor types, mixed with deciduous trees especially in rich types. They are highly diverse, mosaic-like habitats, containing species typical for forests as well as peatland species, because of their forest like conditions and very thin peat layer. Hence, they have significantly higher species richness ^47^ compared to pine mire forests or fens. Poor spruce mire forests are oligotrophic and rich spruce mire forests are meso-eutrophic. Most spruce mires are *Sphagnum* dominated but moving along on the gradient from oligotrophic to eutrophic types, the cover of other mosses and vascular plants, decreasing the cover of *Sphagnums*.

In pine mire forests (Fig. S2c,d; roughly corresponding EUNIS habitat type T3J ^46^) the dominant tree is pine (*Pinus sylvestris*) which may be accompanied by birch (*Betula pubescens*) in rich types. Pine mire forests have typically thick peat layer covered and dominated by peat-forming *Sphagnum* species, accompanied by other mosses. The poor types are ombrotrophic while the rich types vary on a gradient from oligotrophic to mesotrophic sites.

Fens are open peatlands (Fig. S2e,f). The poor fens are ombrotrophic and receive their nutrients mainly from rain water whereas rich fens (category in this study) are oligo-mesotrophic as they receive nutrients also from the surrounding mineral land. Fens are typically *Sphagnum* dominated but when moving along the gradient from ombrotrophic to mesotrophic types, the cover of brown mosses and for example sedges (*Carex* sp.) increases, decreasing the cover of *Sphagnums*. Pine mire forests and fens including high coverage of open water surface were excluded from the site selection.

The sites were chosen to reflect natural variation within each ecosystem type. The selection was based on the Finnish PA network spatial database for habitats and extensive search on old pre- drainage and recent aerial photographs. All sites were visited on site to confirm the observations, and when considered necessary, the peatland type was confirmed based on the occurrence of indicator species, following Økland *et al.* ^28^ . Hydrological independency of the sites was confirmed from topographic data and with field observations. As the study sites are spread along Finland the general environmental conditions (e.g. annual heat sum, average precipitation) vary within the network along a latitudinal gradient. The latitudinal gradient is also reflected as small differences in the vegetation composition among the study sites inside the ecosystem type categories.

Each ecosystem type includes (i) sites that had been drained for forestry during 1960s and 1970s and were restored when included to the monitoring network (‘restored treatment’, approx. 10 sites per ecosystem type), (ii) relatively pristine sites with no drainage (‘pristine control’, approx. 10 sites), and (iii) sites that had been drained and were not restored (‘drained control’, approx. 5 sites) (Fig. S1a, Fig. S2). The treatment, control, and reference sites within each ecosystem type are dispersed in a similar pattern compared to each other. When successful, drainage for forestry results in a decrease of 35-55 cm in the water table level which in turn results in complex changes in nutrient regime and acidity and enhanced tree growth ^48^. Aiming to counteract these changes, restoration was carried out on sites by filling in the ditches and logging the trees that had grown after drainage (if needed) ^49^. Restoration was implemented during autumn-winter between 2007-2014 ^50^.

In restored sites, moss and vascular plant species were monitored prior to restoration (0-year sampling), two (2-year sampling), five (5-year sampling) species and ten (10-year sampling) years after restoration. Similar interval was used in pristine and drained sites, although the time interval between the samplings varies across sites ^50^. The full series of four samplings was completed for 144 sites in total (Fig. S1a). In each site, monitoring was done during the growing season (June-August) on 2007-2022 in ten permanent 1m^2^ plots (Fig. S1b). The plots were systematically situated in two parallel lines, 4m apart from each other. In restored and drained sites, the lines ran parallel to ditches, and the minimum distance to the nearest ditch was 10m. The location of the lines represented typical vegetation of each site, and the location of the first plot was randomized given the criteria above. For all vascular plant and moss species on each plot, a percent cover was visually estimated at the accuracy of one percent. Exceptions were species with a cover of 0.5–0.9% for which a cover of 0.5% was used, and species with a cover of <0.5% for which a cover of 0.2% was used. Species were identified on site, and when not possible, a specimen was taken and later identified under the microscope. Prior to analysis, we removed observations which did not reach species-level resolution, including hybrids. Because identification of particular *Sphagnum* species can be challenging, we grouped these species to avoid uncertainty due to misidentification (Table S1).

### Statistical analysis

We analysed the effect of restoration on individual species occurrences with joint species distribution modelling, using the Hierarchical Modelling of Species Communities framework (HMSC^51,52^). To be able to assess whether the responses to restoration were consistent across ecosystem types, we fitted separate models for each of the ecosystem types. As response variables, we selected the species with at least ten occupancies for poor spruce mire and pine mire forests, as well as poor and rich fens (Table S2). As rich spruce mire forests and pine mire forests showed a higher number of species, we selected for them species with at least twenty occupancies to avoid convergence issues resulting from including a very large number of species. We modelled species occupancy (presence/absence) by a probit model, and conditionally on the presence, cover % (log-transformed, normalized to zero mean and unit variance within each species) with a normal model. Both models had the same structure, as follows. To account for the spatio-temporal structure of the study design where plots are nested within sites which were visited across different years, we included the sampling year, site and plot as random effects. The sampling year and site were included as unstructured random effects, and plots as a spatially explicit random effect. As fixed effects, we included the treatment (a factor with three levels: drained/restored/pristine), time (a continuous variable; 0, 2, 5, 10) and its second order polynomial to allow for unimodal responses, as well as the interaction of treatment and time^2^.

We ran the models with R package Hmsc 3.0.11 ^53^, using the Bayesian framework with Gibbs Markov chain Monte Carlo (MCMC) sampling. We assumed the default prior distributions, with the exception of a1 and a2 parameters for the random effect site which were set both to 100 to increase the shrinkage, and thus avoid modelling noise, in the site-level association matrix. We sampled the posterior distribution with four chains, each for 250 samples with thinning of 1,000, using transient phase of 125,000, and adaptation (the number of MCMC steps at which the adaptation of the number of latent factors is conducted) of 100,000. We evaluated the chain convergence by assessing the effective size of the posterior sample as well as the potential scale reduction factors for each of the estimated parameters (Fig. S3). We assessed model fitting by assessing the difference between the explanatory power (Fig. S4) and predictive power through two-fold cross-validation (Fig. S5) with Tjur’s R^2^ (occupancy) and R^2^ (cover) metrics.

Based on the fitted models, we predicted the abundance of each species (probability of presence × cover given presence) in time for different treatments. From these predictions, we calculated the following five measures informing about different aspects of the effects of restoration on the plant communities.

First, to interpret which species were affected by drainage in the past, we calculated difference in species abundance in pristine versus restored sites at the initial stage of the experiment:

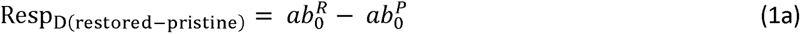

and similarly drained versus restored sites:

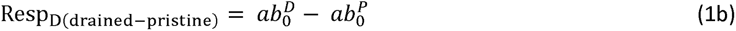

where 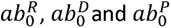 are species’ abundance on plots at time 0 in restored, drained control and pristine control sites, respectively. We calculated the median and the posterior probability for the median being larger than zero, and we considered species having a positive/negative response to drainage if the median for both 1a and 1b were positive/negative with at least 85% posterior probability.

Second, we calculated species response to restoration:

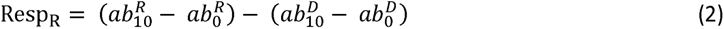

where 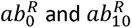 are species’ abundance on plots in restored sites on the time 0 and 10, respectively, and 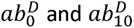 represent corresponding values in drained sites. Resp_R_ takes positive values if abundance change is positive in relation to change in drained sites and negative values if abundance change is negative in relation to change in drained sites.

Third, species abundances in drained and in restored sites may differ as they were not randomly selected. To assess the reliability of inferences of how species respond to restoration, we calculated whether species abundance in the beginning of the experiment differed between the drained and restored sites:

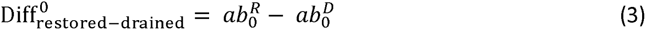

where 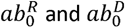 are species’ abundance on plots in drained control and restored sites at time 0, respectively.

Fourth, we calculated whether species abundances were similar in restored and pristine control sites ten years after restoration:

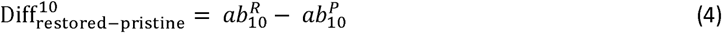

where 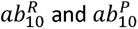 are species’ abundance on plots in restored and pristine control sites at time 10, respectively.

Finally, we calculated whether species abundances change in drained sites:

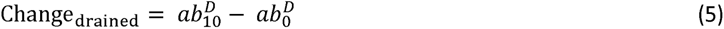

And whether the difference between drained and pristine control sites grew smaller or larger during the study period:

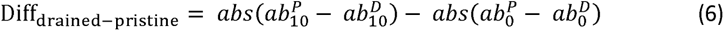

where 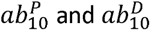 are species’ abundance on plots in pristine and drained control sites at time 10, respectively, and ab^P^ and ab^D^ represent corresponding values at time 0. For all measures from 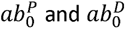 equations (2)-(6), we calculated the median as well as posterior probability for the median being larger than zero. We considered the measure having a high support for the median being positive/negative if the posterior probability is >95% and moderate support if the posterior probability is >80%.

To illustrate variation in community composition among and within ecosystem types and treatments, we produced a model-based ordination using generalized linear latent variable models. The model was fitted with R package gllvm ^54^ for species with at least 10 observations using hurdle beta response model ^55^ and two latent random variables for each site and monitoring year. We assessed model fitting by assessing the difference between the explanatory power and predictive power (based on four-fold cross-validation) (Fig. S9). Further, we calculated a restoration effect for each site based on the latent variables in the ordination, which represent similarities in the species compositions among sites. Restoration effect for each site is the change in average distance to drained sites (time 10 - time 0) minus a change in average distance to pristine sites (that in 10 years, the species composition of time 10 - time 0). Thus, a positive restoration effect means restored site changed more towards pristine sites than drained sites. Conversely, a negative restoration effect means that in 10 years, the species composition of restored site changed more towards drained sites than pristine sites.

## Data availability

The data generated and analysed in this study have been deposited in Zenodo under https://doi.org/10.5281/zenodo.10906943.

## Code availability

All analyses were carried out with the functions and additional packages specified in the methods section in the free and open-source environment R. The specific codes to reproduce the results are presented in Supporting Information 2.

## Acknowledgements

We are deeply thankful for all the researchers, specialists and field workers participating in the development and maintaining the National Peatland Restoration Monitoring Network. We thank Jérémy Cours for the illustrations, Irene Costa Palos for assistance and CSC – IT Center for Science, Finland, for computational resources. The study was funded by the Peatland LIFE (LIFE08 NAT/FIN/000596) and Hydrology LIFE projects (LIFE16 NAT/FIN/000583). In addition, ME was funded by the Finnish Ministry of Environment; OO by the Research Council of Finland (grant no. 336212 and 345110), and the European Union: the European Research Council (ERC) under the European Union’s Horizon 2020 research and innovation programme (grant agreement no. 856506; ERC-synergy project LIFEPLAN); ST by the Research Council of Finland (grant no. 453691); ST and JN by Kone foundation, JSK by the Strategic Research of the Research Council of Finland (grant no. 345710) and NA by the Research Council of Finland (grant no. 342374 and 346492).

## Author contributions

JSK & KA designed the monitoring network, oversaw the maintenance and obtained funding to maintain the monitoring network. ME, OO, JN, NA & ST analysed the data. ME wrote the first draft, and all authors contributed to writing and accepted the final version.

