## Supporting Information 1 for "A large-scale and long-term experiment to identify effectiveness of ecosystem restoration"

### 1. The study set-up

The study set-up includes 151 sites which represent six main ecosystem types (Fig. S1a). Each ecosystem type includes (i) sites that had been drained for forestry during 1960s and 1970s and were restored during the project (‘restored treatment’, approx. 10 sites; Fig. S2), (ii) relatively pristine sites with no drainage (‘pristine control’, approx. 10 sites), and (iii) sites that had been drained and were not restored (‘drained control’, approx. 5 sites). In restored sites, moss and vascular plant species were monitored prior to restoration (0-year sampling), two (2-year sampling), five (5-year sampling) and ten (10-year sampling) years after restoration. Similar interval was used in pristine and drained sites, although the actual number of years between the samplings somewhat differs. The full series of four samplings was completed for 144 sites (Fig. S1a). In each site, species monitoring was done during the growing season (June-August) on 2007-2022 in ten permanent 1m^2^ plots (Fig. S1b). The plots were systematically situated in two parallel lines, 4m apart from each other. In restored and drained sites, the lines ran parallel to ditches, and the minimum distance to the nearest ditch is 10m. The location of the lines represented typical vegetation of each site, and the location of the first plot was randomized given the restrictions above.

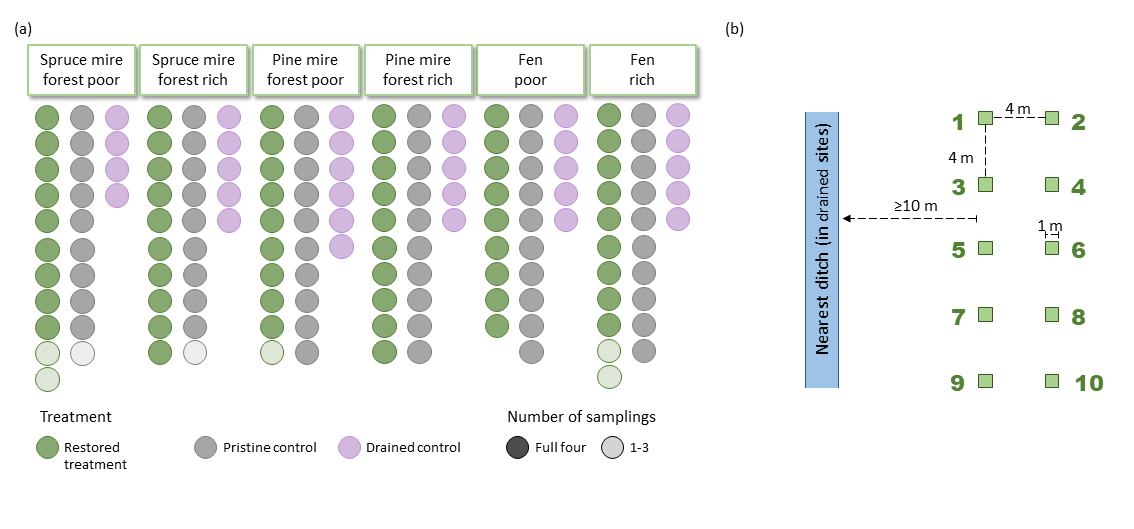

**Fig. S1** The 151 sites included in the study represent six different ecosystem types and three treatments (a). Each site includes ten permanent monitoring plots (b).

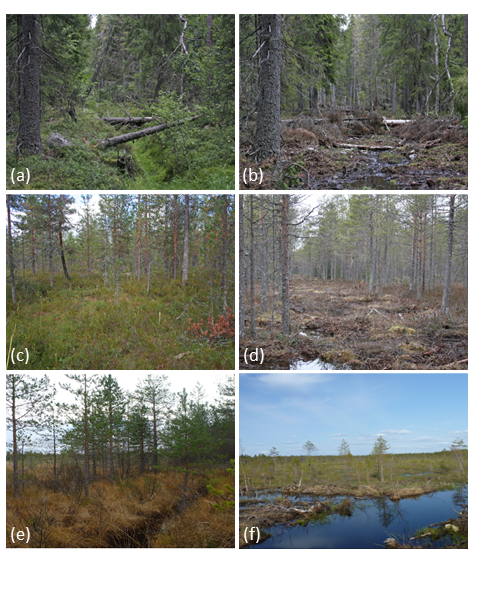

**Fig. S2** A spruce mire forest before (a) and after restoration (b), a pine mire forest before (c) and after restoration (d), and a fen before (e) and after restoration (f). Photos: (a)-(d) Maarit Similä, (e)-(f) Sakari Rehell.

### 2. *Sphagnum* species grouping

Based on extensive consultations of field workers and moss specialists, we identified species groups for each ecosystem type within which species may be easily mixed (Table S1). Misidentification can yield biased inferences on changes in species abundances. For the hmsc analyses, we pooled observations of the species within each group. For the model-based ordination, ecosystem types were modeled simultaneously. Hence, we used a different approach. For each site separately, we observed which of the species belonging to group have been identified. We assumed that the correct species have been identified at least one of the sampling times. Then, we calculated the total cover of the group for each sample (site_plot_time) and divided this equally between all species belonging to the group which had been observed during the sampling.

**Table S1** Grouping of specific *Sphagnum* species in different ecosystem types.

| **Ecosystem type** | **Group** | **Species included** |
| --- | --- | --- |
| Spruce mire forest poor | Spha I | *S. girgensohnii, S. riparium, S. recurvum coll., S. russowii* |
| Spruce mire forest rich | Spha II | *S. girgensohnii, S. riparium, S. russowii* |
| Pine mire forest poor | Spha III | *S. recurvum coll., S. capillifolium, S. balticum* |
| Pine mire forest rich | Spha IV | *S. recurvum coll., S. balticum* |
| Fen poor | Spha IV | *S. recurvum coll., S. balticum* |
| Fen rich | No grouping |  |

### 3. Hmsc modelling: convergence, explanatory and predictive power

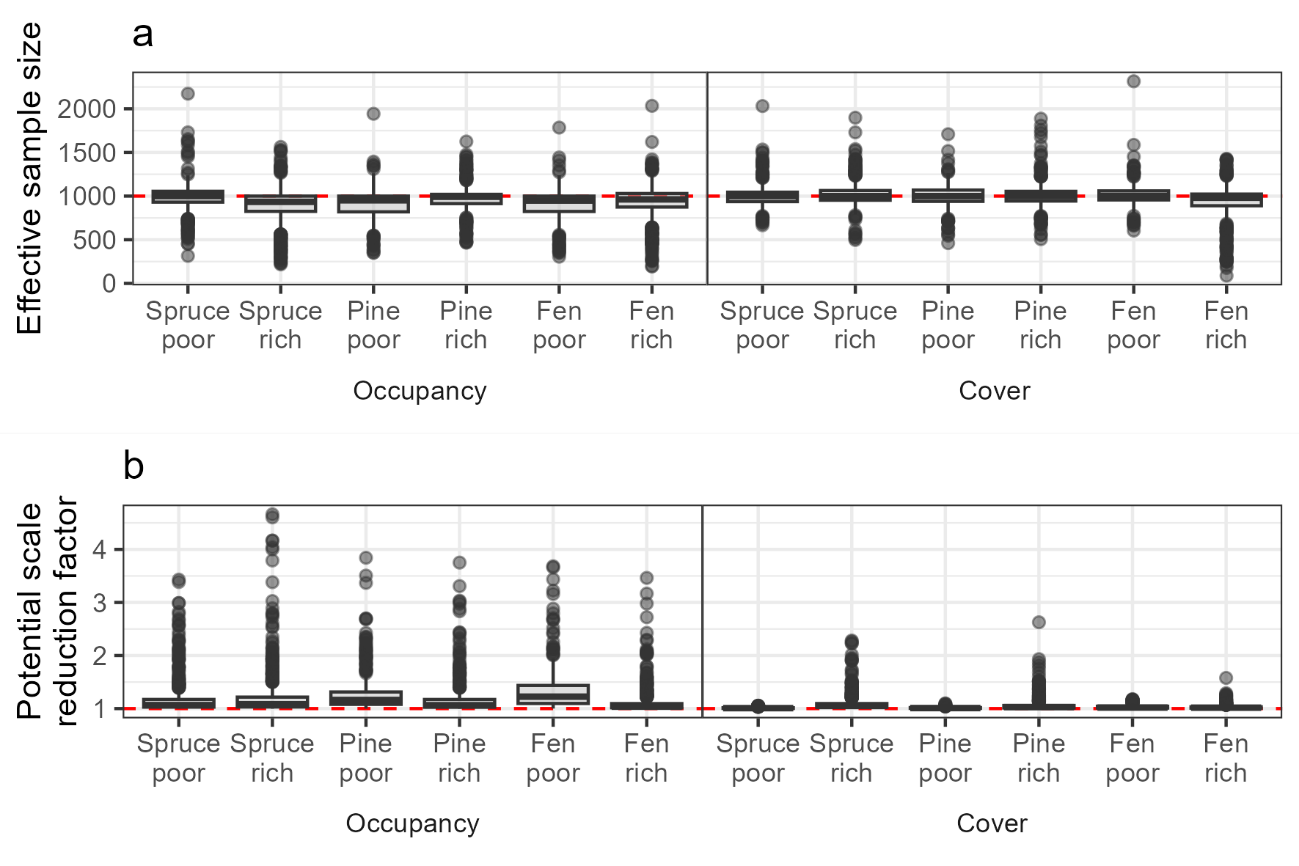

**Fig. S3** We evaluated the convergence of beta parameters with the effective size of the posterior sample (a) and potential scale reduction factor (b). Convergence was good for cover models but satisfactory for occupancy models. However, non-ideal convergence of non-normal models is found often when applying HMSC as well as other approaches using posterior sampling with MCMC of large, multi-dimensional data (see Tikhonov *et al.* 2020 Methods in Ecology and Evolution 11: 442-447, as referred in the main text).

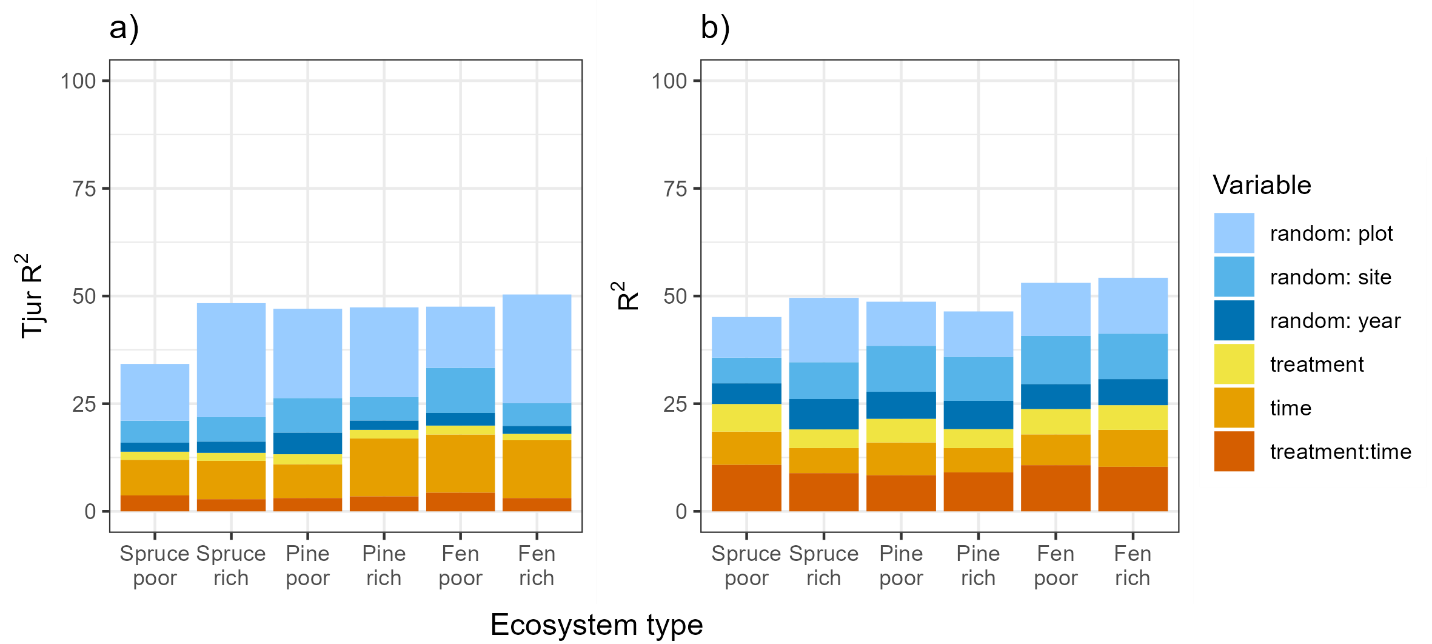

**Fig. S4** The mean explanatory power for species occupancy models (Tjur’s R^2^) (a) and cover models (R^2^) (b).

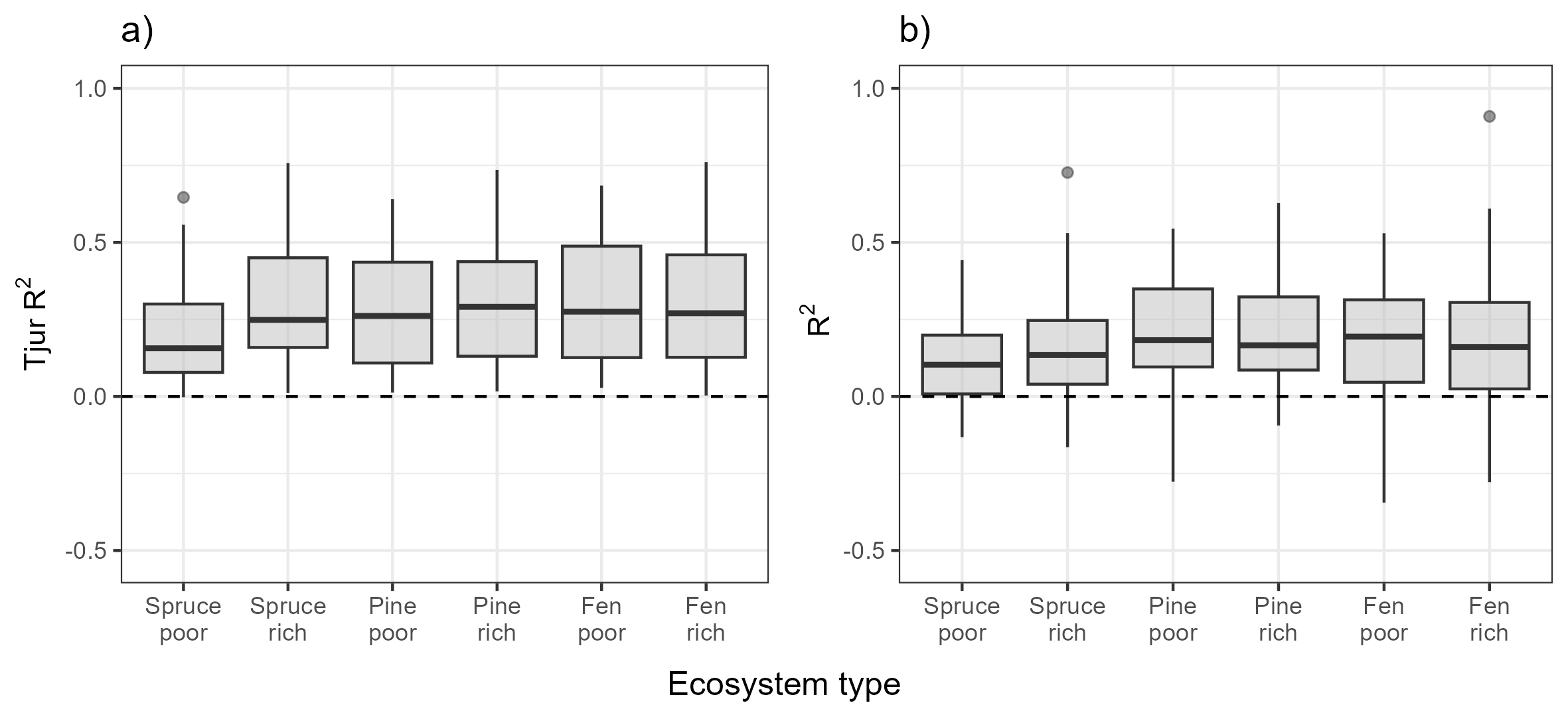

**Fig. S5** The predictive power for occupancy (a) and cover (b) models for different ecosystem types. The median is shown as the black line, and lower and upper hinges of the box correspond to the 25th and 75th percentiles. The whiskers extend from the hinge to the largest/smallest value no further than 1.5 × inter-quartile range from the hinge (where IQR is the or distance between the first and third quartiles). Points beyond the end of the whiskers are plotted individual

### 4. Species and their responses

**Table S2** Species included in the analysis. Species shortening, scientific name, group (moss or vascular plant, ‘Vasc’), and their occurrence in the analysis of each ecosystem type.

| **Species shortening** | **Species scientific name** | **Group** | **Spruce mire forest** | | **Pine mire forest** | | **Fen** | |
| --- | --- | --- | --- | --- | --- | --- | --- | --- |
|  |  |  | **Poor** | **Rich** | **Poor** | **Rich** | **Poor** | **Rich** |
| Andr poli | *Andromeda polifolia* | Vasc |  | x | x | x | x | x |
| Athy fili | *Athyrium filix-femina* | Vasc |  | x |  |  |  |  |
| Aula palu | *Aulacomnium palustre* | Moss | x | x | x | x | x | x |
| Aven flex | *Avenella flexuosa* | Vasc | x | x |  |  |  |  |
| Betu nana | *Betula nana* | Vasc |  |  | x | x | x | x |
| Betu pube | *Betula pubescens* | Vasc | x | x | x | x |  | x |
| Brac sale | *Brachythecium salebrosum* | Moss |  | x |  |  |  |  |
| Cala arun | *Calamagrostis arundinacea* | Vasc | x |  |  |  |  |  |
| Cala cane | *Calamagrostis canescens* | Vasc |  | x |  |  |  |  |
| Cala phra | *Calamagrostis phragmitoides* | Vasc |  | x |  |  |  |  |
| Call cord | *Calliergon cordifolium* | Moss |  | x |  |  |  |  |
| Call vulg | *Calluna vulgaris* | Vasc |  |  | x | x | x | x |
| Care aqua | *Carex aquatilis* | Vasc |  |  |  | x |  |  |
| Care brun | *Carex brunnescens* | Vasc |  | x |  |  |  |  |
| Care cane | *Carex canescens* | Vasc | x | x |  | x |  | x |
| Care cesp | *Carex cespitosa* | Vasc |  | x |  |  |  |  |
| Care chor | *Carex chordorrhiza* | Vasc |  | x |  | x |  | x |
| Care dioi | *Carex dioica* | Vasc |  |  |  | x |  | x |
| Care disp | *Carex disperma* | Vasc | x | x |  |  |  |  |
| Care echi | *Carex echinata* | Vasc |  | x |  | x |  | x |
| Care glob | *Carex globularis* | Vasc | x | x | x | x |  |  |
| Care lasi | *Carex lasiocarpa* | Vasc |  | x |  | x |  | x |
| Care limo | *Carex limosa* | Vasc |  |  |  | x | x | x |
| Care loli | *Carex loliacea* | Vasc | x |  |  |  |  |  |
| Care nigr | *Carex nigra* | Vasc |  |  |  | x |  |  |
| Care pauc | *Carex pauciflora* | Vasc |  |  | x | x | x | x |
| Care paup | *Carex paupercula* | Vasc |  |  |  | x |  | x |
| Care rost | *Carex rostrata* | Vasc |  |  |  | x |  | x |
| Care vagi | *Carex vaginata* | Vasc |  | x |  |  |  |  |
| Care vesi | *Carex vesicaria* | Vasc |  | x |  |  |  |  |
| Cham caly | *Chamaedaphne calyculata* | Vasc | x | x | x | x | x | x |
| Coma palu | *Comarum palustre* | Vasc |  | x |  | x |  | x |
| Dact macu | *Dactylorhiza maculata* | Vasc | x |  |  |  |  | x |
| Desc cesp | *Deschampsia cespitosa* | Vasc |  | x |  |  |  |  |
| Dicr maju | *Dicranum majus* | Moss | x | x |  | x |  |  |
| Dicr poly | *Dicranum polysetum* | Moss | x | x | x | x | x | x |
| Dicr scop | *Dicranum scoparium* | Moss | x | x | x | x | x | x |
| Dicr undu | *Dicranum undulatum* | Moss |  |  | x |  | x | x |
| Dros angl | *Drosera anglica* | Vasc |  |  |  |  | x | x |
| Dros rotu | *Drosera rotundifolia* | Vasc |  |  | x | x | x | x |
| Dryo cart | *Dryopteris carthusiana* | Vasc | x | x |  |  |  | x |
| Dryo expa | *Dryopteris expansa* | Vasc |  | x |  |  |  |  |
| Empe nigr | *Empetrum nigrum* | Vasc | x |  | x | x | x | x |
| Equi arve | *Equisetum arvense* | Vasc |  | x |  |  |  |  |
| Equi fluv | *Equisetum fluviatile* | Vasc |  | x |  | x |  | x |
| Equi palu | *Equisetum palustre* | Vasc |  | x |  | x |  |  |
| Equi prat | *Equisetum pratense* | Vasc |  | x |  |  |  |  |
| Equi sylv | *Equisetum sylvaticum* | Vasc | x | x |  |  |  |  |
| Erio angu | *Eriophorum angustifolium* | Vasc |  |  |  | x | x | x |
| Erio vagi | *Eriophorum vaginatum* | Vasc | x | x | x | x | x | x |
| Fili ulma | *Filipendula ulmaria* | Vasc |  | x |  |  |  |  |
| Fran alnu | *Frangula alnus* | Vasc | x |  |  |  |  |  |
| Gera sylv | *Geranium sylvaticum* | Vasc |  | x |  |  |  |  |
| Good repe | *Goodyera repens* | Vasc |  | x |  |  |  |  |
| Gymn dryo | *Gymnocarpium dryopteris* | Vasc |  | x |  |  |  |  |
| Helo blan | *Helodium blandowii* | Moss |  |  |  |  |  | x |
| Hylo sple | *Hylocomium splendens* | Moss | x | x | x | x |  |  |
| Hylo triq | *Hylocomiadelphus triquetrus* | Moss |  | x |  |  |  |  |
| Linn bore | *Linnaea borealis* | Vasc | x | x |  |  |  |  |
| Loes badi | *Loeskypnum badium* | Moss |  |  |  |  |  | x |
| Luzu pilo | *Luzula pilosa* | Vasc | x | x |  |  |  |  |
| Lysi euro | *Lysimachia europaea* | Vasc | x | x |  |  |  | x |
| Lysi thyr | *Lysimachia thyrsiflora* | Vasc |  | x |  |  |  |  |
| Lysi vulg | *Lysimachia vulgaris* | Vasc |  | x |  |  |  |  |
| Maia bifo | *Maianthemum bifolium* | Vasc | x | x | x |  |  |  |
| Mela prat | *Melampyrum pratense* | Vasc | x | x | x | x | x | x |
| Mela sylv | *Melampyrum sylvaticum* | Vasc | x |  |  |  |  |  |
| Meny trif | *Menyanthes trifoliata* | Vasc |  | x |  | x |  | x |
| Moli caer | *Molinia caerulea* | Vasc |  | x |  | x |  | x |
| Neot cord | *Neottia cordata* | Vasc | x |  |  |  |  |  |
| Orth secu | *Orthilia secunda* | Vasc | x | x |  |  |  |  |
| Oxal acet | *Oxalis acetosella* | Vasc | x | x |  |  |  |  |
| Pari quad | *Paris quadrifolia* | Vasc |  | x |  |  |  |  |
| Pheg conn | *Phegopteris connectilis* | Vasc |  | x |  |  |  |  |
| Pice abie | *Picea abies* | Vasc | x | x | x | x |  | x |
| Pinu sylv | *Pinus sylvestris* | Vasc | x |  | x | x | x | x |
| Plag curv | *Plagiothecium curvifolium* | Moss | x | x |  |  |  |  |
| Plag cusp | *Plagiomnium cuspidatum* | Moss |  |  | x |  |  |  |
| Plag dent | *Plagiothecium denticulatum* | Moss | x | x |  |  |  |  |
| Plag elli | *Plagiomnium ellipticum* | Moss |  | x |  |  |  |  |
| Plag laet | *Plagiothecium laetum* | Moss | x | x |  |  |  |  |
| Pleu schr | *Pleurozium schreberi* | Moss | x | x | x | x | x | x |
| Pohl nuta | *Pohlia nutans* | Moss | x | x | x | x | x | x |
| Pohl spha | *Pohlia sphagnicola* | Moss |  |  | x |  |  | x |
| Poly comm | *Polytrichum commune* | Moss | x | x |  | x |  | x |
| Poly form | *Polytrichum formosum* | Moss | x |  |  |  |  |  |
| Poly juni | *Polytrichum juniperinum* | Moss |  |  | x |  |  |  |
| Poly long | *Polytrichum longisetum* | Moss | x | x |  | x |  |  |
| Poly stri | *Polytrichum strictum* | Moss |  |  | x | x | x | x |
| Poly swar | *Polytrichum swartzii* | Moss |  |  |  |  |  | x |
| Pseu cinc | *Pseudobryum cinclidioides* | Moss |  | x |  |  |  |  |
| Ptil cris | *Ptilium crista-castrensis* | Moss | x |  |  |  |  | x |
| Pyro mino | *Pyrola minor* | Vasc |  | x |  |  |  |  |
| Ranu repe | *Ranunculus repens* | Vasc |  | x |  |  |  |  |
| Rhiz pseu | *Rhizomnium pseudopunctatum* | Moss |  | x |  |  |  |  |
| Rhod rose | *Rhodobryum roseum* | Moss |  | x |  |  |  |  |
| Rhod tome | *Rhododendron tomentosum* | Vasc |  |  | x | x |  | x |
| Rhyn alba | *Rhynchospora alba* | Vasc |  |  |  |  | x |  |
| Rubu arct | *Rubus arcticus* | Vasc |  | x |  |  |  |  |
| Rubu cham | *Rubus chamaemorus* | Vasc | x | x | x | x | x | x |
| Rubu idae | *Rubus idaeus* | Vasc |  | x |  |  |  |  |
| Rubu saxa | *Rubus saxatilis* | Vasc |  | x |  |  |  |  |
| Sali lapp | *Salix lapponum* | Vasc |  |  |  |  |  | x |
| Sali myrt | *Salix myrtilloides* | Vasc |  |  |  | x |  | x |
| Sani unci | *Sanionia uncinata* | Moss | x | x |  |  |  |  |
| Sarm exan | *Sarmentypnum exannulatum* | Moss |  |  |  |  |  | x |
| Sche palu | *Scheuchzeria palustris* | Vasc |  |  |  | x | x | x |
| Sciu curt | *Sciuro-hypnum curtum* | Moss | x | x |  |  |  |  |
| Sciu refl | *Sciuro-hypnum reflexum* | Moss | x | x |  |  |  |  |
| Sciu star | *Sciuro-hypnum starkei* | Moss | x | x |  |  |  |  |
| Sorb aucu | *Sorbus aucuparia* | Vasc | x | x |  |  |  |  |
| Spha annu/jens | *Sphagnum annulatum/jensenii* | Moss |  |  |  | x |  | x |
| Spha aong | *Sphagnum aongstroemii* | Moss |  |  |  |  |  | x |
| Spha balt | *Sphagnum balticum* | Moss |  |  | gIII | gIV | gIV | x |
| Spha capi | *Sphagnum capillifolium* | Moss | x | x | gIII | x | x | x |
| Spha cent | *Sphagnum centrale* | Moss | x | x | x |  |  |  |
| Spha comp | *Sphagnum compactum* | Moss |  |  |  |  | x | x |
| Spha cusp | *Sphagnum cuspidatum* | Moss |  |  |  |  | x |  |
| Spha fimb | *Sphagnum fimbriatum* | Moss | x |  |  |  |  |  |
| Spha fusc | *Sphagnum fuscum* | Moss |  |  | x | x | x | x |
| Spha girg | *Sphagnum girgensohnii* | Moss | gI | gII |  |  |  |  |
| Spha lind | *Sphagnum lindbergii* | Moss |  |  |  |  |  | x |
| Spha maju | *Sphagnum majus* | Moss |  |  |  |  | x | x |
| Spha medi coll. | *Sphagnum medium coll.* | Moss | x | x | x | x | x | x |
| Spha obtu | *Sphagnum obtusum* | Moss |  |  |  |  |  | x |
| Spha papi | *Sphagnum papillosum* | Moss |  |  | x | x | x | x |
| Spha pulc | *Sphagnum pulchrum* | Moss |  |  |  |  |  | x |
| Spha quin | *Sphagnum quinquefarium* | Moss |  | x |  |  |  |  |
| Spha recu coll. | *Sphagum recurvum coll.* | Moss | gI | x | gIII | gIV | gIV | x |
| Spha ripa | *Sphagnum riparium* | Moss | gI | gII |  | x |  | x |
| Spha rube/warn | *Sphagnum rubellum/warnstorfii* | Moss |  | x | x | x | x | x |
| Spha russ | *Sphagnum russowii* | Moss | gI | gII | x | x | x | x |
| Spha squa | *Sphagnum squarrosum* | Moss | x | x |  |  |  |  |
| Spha subf/subn | *Sphagnum subfulvum/subnites* | Moss |  |  |  |  |  | x |
| Spha subs | *Sphagnum subsecundum* | Moss |  |  |  |  |  | x |
| Spha tene | *Sphagnum tenellum* | Moss |  |  |  |  | x | x |
| Spha tere | *Sphagnum teres* | Moss |  |  |  |  |  | x |
| Spha wulf | *Sphagnum wulfianum* | Moss | x |  |  |  |  |  |
| Spin anno | *Spinulum annotinum* | Vasc |  | x |  |  |  |  |
| Stra stra | *Straminergon stramineum* | Moss | x | x | x | x | x | x |
| Tetr pell | *Tetraphis pellucida* | Moss | x | x |  |  |  |  |
| Tric alpi | *Trichophorum alpinum* | Vasc |  |  |  |  |  | x |
| Tric cesp | *Trichophorum cespitosum* | Vasc |  |  |  |  | x | x |
| Vacc micr | *Vaccinium microcarpum* | Vasc |  |  | x | x | x | x |
| Vacc myrt | *Vaccinium myrtillus* | Vasc | x | x | x | x | x | x |
| Vacc oxyc | *Vaccinium oxycoccos* | Vasc | x | x | x | x | x | x |
| Vacc ulig | *Vaccinium uliginosum* | Vasc | x | x | x | x | x | x |
| Vacc viti | *Vaccinium vitis-idaea* | Vasc | x | x | x | x | x | x |
| Viol epip | *Viola epipsila* | Vasc |  | x |  |  |  |  |
| Viol palu | *Viola palustris* | Vasc |  | x |  |  |  |  |
| Warn flui | *Warnstorfia fluitans* | Moss |  |  |  |  | x | x |

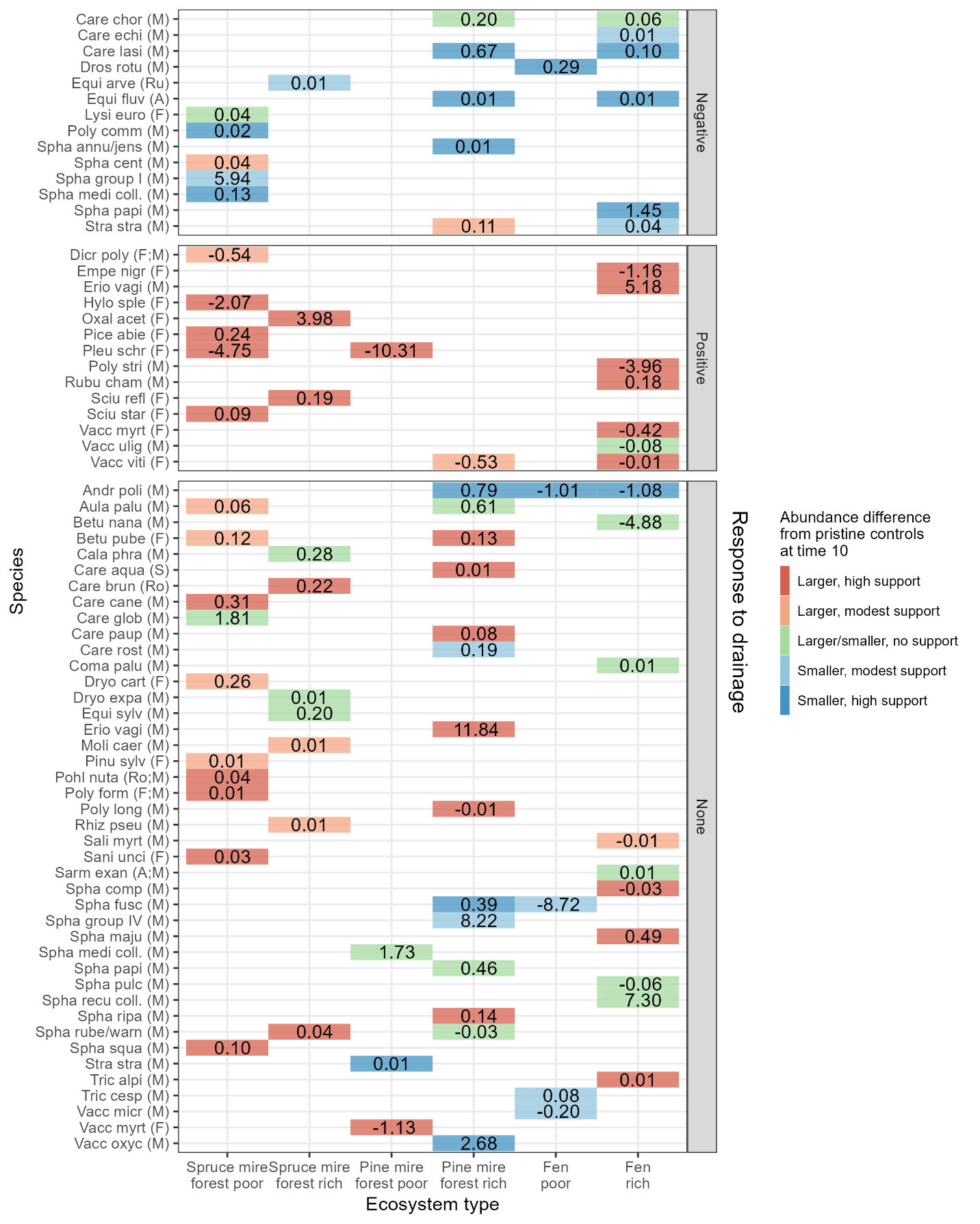

**Fig. S6** For each species responding to restoration (as in Fig. 4), it is indicated by the color whether their abundance differs at restored sites and control sites ten years after restoration. If the abundance does not differ (posterior probability <80%), species is shown in green. If the abundance is larger in restored than in pristine control with high support (>95%) or moderate support (>80%) the species in dark and light red, respectively. If the abundance is smaller in restored than in pristine control with high support (>95%) or moderate support (>80%) the species in dark and light blue respectively. In addition, median of the response is shown, separately according to their response to drainage in the given ecosystem type. The species are separated according to their response to drainage. Species that had had negative response to drainage, have largely increased to similar level of abundances than in pristine controls. Species that had had positive response to drainage, have decreased but have still larger abundances than pristine controls. Species that had no response to drainage often increase to have larger abundances than in pristine controls. Species’ full names can be found from Table S2. Species primary habitat type is shown in parentheses [F = forests, M = mires (including mire forests and fens), A = aquatic habitats, S = shores, Ro = rock outcrops and boulder fields, Ru = rural biotopes and cultural habitats]

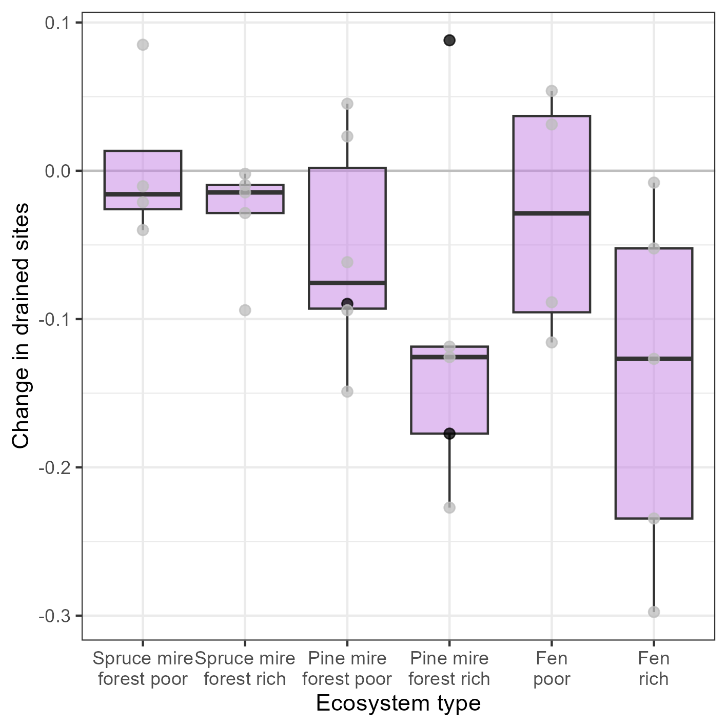

**Fig. S7** The change in drained sites (based on the latent variables from the model-based ordination in Fig. 2) for each drained site is positive if species composition in a site changed towards pristine sites from before restoration to ten years after restoration. Conversely, the measure is negative if species composition changed increasingly dissimilar to pristine sites. The thick line corresponds median, and lower and upper hinges to the 25^th^ and 75^th^ percentiles. The upper/lower whisker extends to the largest/smallest value that is at maximum 1.5 × distance between the 25^th^ and 75^th^ percentiles. Data are shown as points, and the black points are for those three sites for which the ditches have been re-opened.

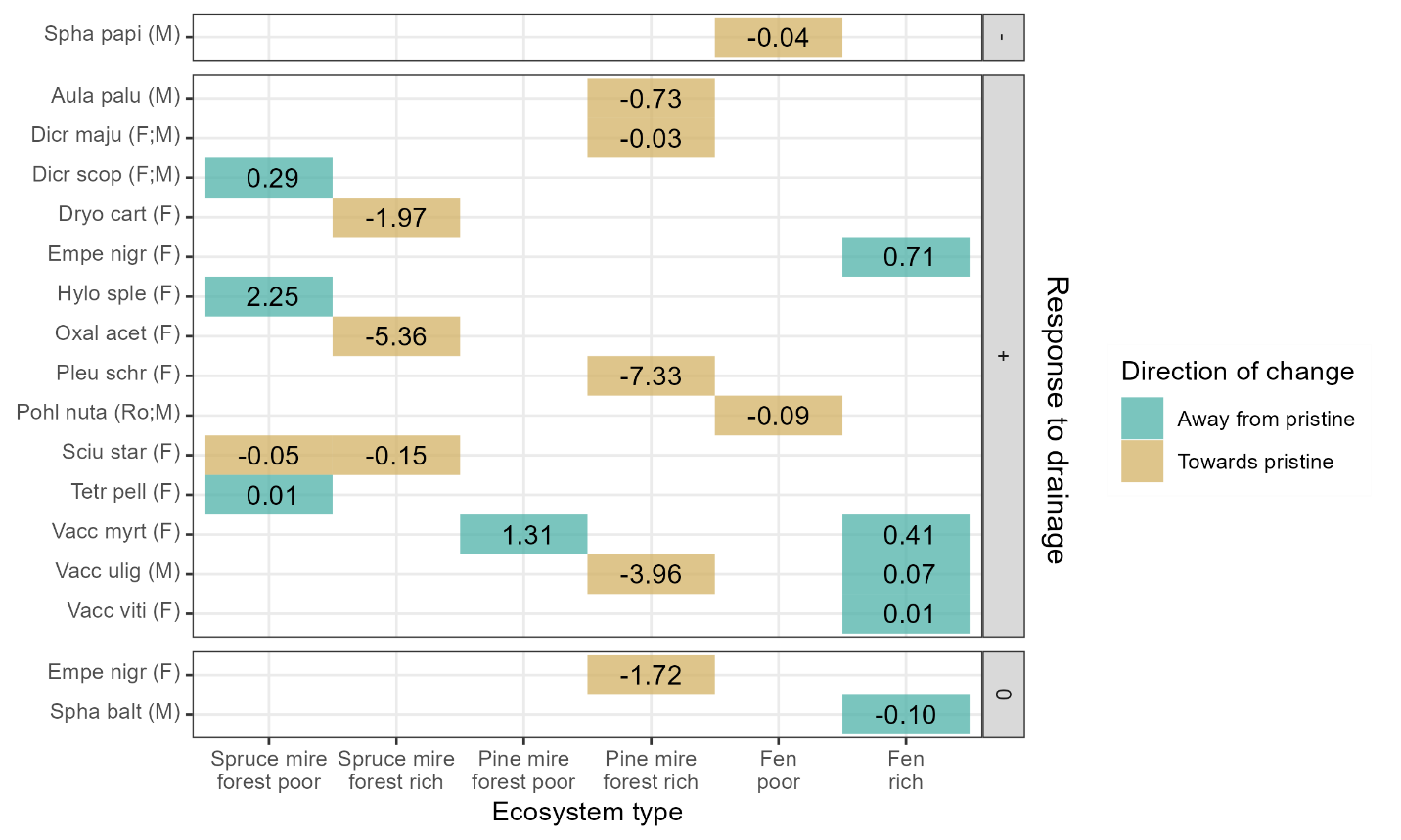

**Fig. S8** For each species chancing in drained sites with high statistical support (95% posterior probability) and for which the direction of change in species abundance is away from or towards that in pristine controls with high statistical support, median of the response is shown, separately according to their response to drainage in the given ecosystem type. The color indicates whether the direction of change in species abundance is away from (green) or towards (yellow) that in pristine sites. Only response median values larger (smaller) than 0.01 (-0.01) are shown. Species’ full names can be found from Table S2. Species primary habitat type is shown in parentheses [F = forests, M = mires (including mire forests and fens), A = aquatic habitats, S = shores, Ro = rock outcrops and boulder fields, Ru = rural biotopes and cultural habitats]

### 5. Model-based ordination: explanatory and predictive power

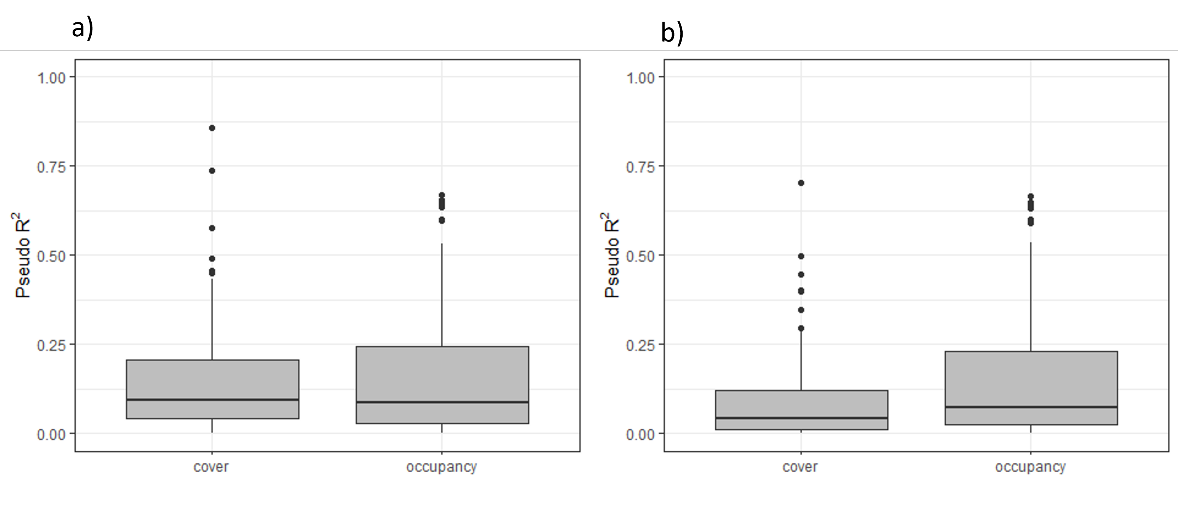

**Fig. S8** Species-specific explanatory power (a) and predictive power (based on 4-fold cross-validation) (b) for the ordination model. The pseudo-R^2^ measures were Tjur’s R^2^ for the binomial probit occupancy model and a pseudo-R^2^ by Ferrari and Cribari-Neto ^1^ for the beta cover model.

^1^ Ferrari S. & Cribari-Neto F. (2004) Beta regression for modelling rates and proportions. Journal of Applied Statistics 31:799-815.
