## Supporting Information 2 for "A large-scale and long-term experiment to identify effectiveness of ecosystem restoration"

### 1. HMSC analyses

#### 1.1 Data preparation

Prior to analysis, we have pooled the observations of specific Sphagnum species (see Table S1 in Supporting Information 1).

```
library(readr)
library(dplyr)
library(readxl)

#load the species data
load("Data_Napalm_FINAL.RData")
da = DATA
da1 = data.frame(site = as.factor(paste0("site",da$ID)),
                 plot = as.factor(paste0("site",da$ID,"_plot",da$Plot)),
                 year = da$Monitor_year,
                 time = as.factor(da$Time),
                 plot.time = as.factor(paste0("site",da$ID,"_plot",
                                              da$Plot,"_time",da$Time)),
                 group = as.factor(da$Group),
                 species = as.factor(da$Species),
                 cover = da$Cover)
```

The data `da1` includes 'site' (a unique number for a site), 'plot' (a study plot within a site: 1-10), 'year' (the year for the sampling), 'time' (the sampling time: 0/2/5/10 years after restoration), 'plot.time' (specifying each sample). 'group' (1=vascular plant, 2=moss), 'species' (a shortening for each species), 'cover' (% cover in a plot)

```
##    site        plot year time         plot.time group   species cover
## 1 site1 site1_plot2 2009    0 site1_plot2_time0     1 Dryo cart   0.2
## 2 site1 site1_plot4 2009    0 site1_plot4_time0     1 Dryo cart   3.0
## 3 site1 site1_plot5 2009    0 site1_plot5_time0     1 Dryo cart   4.0
## 4 site1 site1_plot1 2009    0 site1_plot1_time0     1 Maia bifo   3.0
## 5 site1 site1_plot2 2009    0 site1_plot2_time0     1 Maia bifo   3.0
## 6 site1 site1_plot3 2009    0 site1_plot3_time0     1 Maia bifo   2.0
```

We start by making the trait matrix `TrData.i` specifying to which group (vascular plant/moss) each species belongs.

```
ta = table(da1$species) #occurrence for all species
spp = names(ta) #names of the species
nsp = length(spp) #number of species

ind.sp = match(da1$species,spp)
temp = rep(0,nsp)
for(i in 1:nsp){
  temp[i] = median(as.numeric(da1$group[ind.sp==i]))
}

Tr = rep("NA",nsp)
Tr[temp==1] = "vascular.plant"
Tr[temp==2] = "moss"

TrData.i = data.frame(group = as.factor(Tr))
rownames(TrData.i) = spp
```

```
##                      group
## Agro cani vascular.plant
## Agro capi vascular.plant
## Alnu glut vascular.plant
## Alnu inca vascular.plant
## Ambl serp           moss
## Andr poli vascular.plant
```

Next, we make the species matrix `Y.i.AL` with species as columns and sampling units 'plot.time' as rows.

```
sampling.units = levels(da1$plot.time)
ny = length(sampling.units)
ind.su = match(da1$plot.time,sampling.units)

Y.i.AL = matrix(0,nrow = ny, ncol = nsp)
for(i in 1:length(ind.su)){
  Y.i.AL[ind.su[i],ind.sp[i]] = da1$cover[i]
}
colnames(Y.i.AL) = spp
rownames(Y.i.AL) = sampling.units
```

```
##                      group
## Agro cani vascular.plant
## Agro capi vascular.plant
## Alnu glut vascular.plant
## Alnu inca vascular.plant
## Ambl serp           moss
## Andr poli vascular.plant
```

Then we read in the site-level explanatory variables.

```
meta <- read_excel("meta.xlsx")
ta2 = data.frame(site = as.factor(paste0("site",meta$ID)),
                 treatment = as.factor(meta$Treatment),
                 type = as.factor(meta$Type),
                 x = as.numeric(meta$x),
                 y = as.numeric(meta$y))
```

The data includes 'site' (similar to `Y.i.AL`), 'treatment' (restored / pristine / drained), 'type' (Spruce.P / Spruce.R / Pine.P / Pine.R / Fen.P / Fen.R) and coordinates (x,y) for each site.

```
##    site treatment     type        x        y
## 1 site1  restored Spruce.P 25.19755 60.28960
## 2 site2  restored Spruce.P 23.06104 60.23510
## 3 site3  restored Spruce.P 25.16245 61.30178
## 4 site4  restored Spruce.P 28.65656 62.48171
## 5 site5  restored Spruce.P 28.80704 62.44713
## 6 site6  restored Spruce.P 25.56754 62.21680
```

We continue by making the spatial matrix `sitexy` and the environmental data matrix `Xdata`.

```
#coordinates
sitexy = as.matrix(cbind(ta2$x,ta2$y))
colnames(sitexy) = c("x","y")
rownames(sitexy) = ta2$site

#explanatory variables
si = rep(0,ny) #site
pl = rep(0,ny) #plot
ye = rep(0,ny) #year
ti = rep(0,ny) #time

for(i in 1:ny){
  da3sel = da1[ind.su==i,]
  si[i] = da3sel[1,]$site
  pl[i] = da3sel[1,]$plot
  ye[i] = da3sel[1,]$year
  ti[i] = da3sel[1,]$time
}

XData = data.frame(sample = as.factor(sampling.units),
                   site = as.factor(levels(da1$site)[si]),
                   plot = as.factor(levels(da1$plot)[pl]),
                   year = ye,
                   time = as.numeric(as.character(levels(da1$time)[ti])))

ind = match(XData$site,ta2$site)

XData$treatment = as.factor(ta2[ind,]$treatment)
XData$type = as.factor(ta2[ind,]$type)
```

The data is analysed separately for each ecosystem type (Spruce.P/Spruce.R/Pine.P/Pine.R/Fen.P/Fen.R), and the following procedure is looped over the six types. First, we divide environmental data `XData` and spatial data `sitexy`.

```
#environmental data (XData)
XData.sub = subset(XData, type== 'Spruce.P')
XData.sub = droplevels(XData.sub)
rownames(XData.sub) <- c(1:nrow(XData.sub))

#spatial data (sitexy)
sub.sites <- XData.sub$site
sitexy.tmp <- data.frame(sitexy)
sitexy.tmp$site <- NA
sitexy.tmp$site <- row.names(sitexy)

sitexy.sub = sitexy.tmp[sitexy.tmp$site %in% sub.sites,]
sitexy.sub = droplevels(sitexy.sub[,1:2])
```

Next, we divide `Y.i.AL` to two Y-matrices: `Y.ipa` presence/absence of the species with occurrence > limit, and 2) abundance of the species with occurrence > limit. In addition, we make a trait matrices.

```
#set the limit for minimum occurrence for a species
limit <- 20 # for rich spruce mire forests and rich pine mire forests
limit <- 10 #other types

sub.samples <- XData.sub$sample #samples in a given type

Y.i.AL.tmp <- data.frame(Y.i.AL) #all species, all types
Y.i.AL.tmp$sample <- NA
Y.i.AL.tmp$sample <- rownames(Y.i.AL) #set samples as one column

Y.i.tmp = Y.i.AL.tmp[Y.i.AL.tmp$sample %in% sub.samples,] #samples for the type
Y.i.tmp2 = Y.i.tmp[,1:(ncol(Y.i.tmp)-1)]
Y.i.tmp2 = droplevels(Y.i.tmp2)
Y.i.tmp2 = as.matrix(Y.i.tmp2)
colnames(Y.i.tmp2) = colnames(Y.i.AL)

Y.i = Y.i.tmp2
Y.i[Y.i > 0] <- 1 #from abundance to P/A

#species occurring more often than the limit
cS = colSums(Y.i)
spp.c = names(cS[cS>=limit])

Y.ipa = Y.i[,colnames(Y.i) %in% spp.c] #occurrence
Y.iab.tmp = Y.i.tmp2[,colnames(Y.i.tmp2) %in% spp.c]
Y.iab.tmp[Y.iab.tmp==0] = NA #no occurrence -> NA
Y.iab = scale(log(Y.iab.tmp)) #log-transform and scale abundance

#traits
TrData.sub = data.frame(TrData.i[rownames(TrData.i) %in% spp.c,])
colnames(TrData.sub) <- c('group')
rownames(TrData.sub) <- spp.c
```

In the end, we have four matrices for each type: 1) `XData.sub` environmental variables 2) `sitexy.sub` spatial data 3) `Y.ipa` species presence/absence 4) `Y.iab` species abundance, given presence)

#### 1.2 Models

We define two models for each ecosystem type.

```
library(Hmsc)

#study design
studyDesign = data.frame(site = XData.sub$site,
                         plot = XData.sub$plot,
                         sample = XData.sub$sample,
                         year = as.factor(XData.sub$year))

#random levels
rL.site = HmscRandomLevel(sData = sitexy.sub)
rL.plot = HmscRandomLevel(units = levels(studyDesign$plot))
rL.year = HmscRandomLevel(units = levels(studyDesign$year))

#set a1 and a2 priors for random factor = site
rL.site = setPriors(rL.site, a1=100,a2=100)

#remove information not needed from XData
XData.sub$site = NULL
XData.subplot = NULL
XData.sub$sample = NULL
XData.sub$year = NULL

#formula for "environmental" effects
XFormula = ~ poly(time,degree=2,raw=TRUE)*treatment

#formula for traits
TrFormula = ~ group

#number of species
ns = dim(Y.ipa)[2]

#model for presence/absence
m1 = Hmsc(Y=Y.ipa, XData = XData.sub,  XFormula = XFormula,
          studyDesign=studyDesign,
          TrData = TrData.sub, TrFormula = TrFormula,
          ranLevels={list(site = rL.site,
                          plot = rL.plot,
                          year = rL.year)},
          distr = c(rep("probit",ns)))

#model for abundance, given presence
m2 = Hmsc(Y=Y.iab, XData = XData.sub,  XFormula = XFormula,
          studyDesign=studyDesign,
          TrData = TrData.sub, TrFormula = TrFormula,
          ranLevels={list(site = rL.site,
                          plot = rL.plot,
                          year = rL.year)},
          distr = c(rep("normal",ns)))
```

#### 1.3 Model fitting

```
nChains=4
samples=250
thin=1000

m = sampleMcmc(m, thin=thin, samples=samples, nChains=nChains,
               transient=500*thin)
```

#### 1.4 Model convergence

```
library(Hmsc)

#all models as a list
load("models_thin_1000_samples_250_chains_4.RData")

ess_list = list()
gel_list = list()

for(i in 1:12){

  m =  models[[i]]
  mpost = convertToCodaObject(m)

  #Effective sample size
  efs <- data.frame(effectiveSize(mpost$Beta))
  efs$model.no <- paste(i)

  ess_list[[i]] = efs

  #Potential scale reduction factor
  gel <- data.frame(gelman.diag(mpost$Beta, multivariate=FALSE)$psrf)
  gel$model.no <- paste(i)

  gel_list[[i]] = gel
}
```

#### 1.5 Variance partition

```
library(Hmsc)

#all models as a list
load("models_thin_1000_samples_250_chains_4.RData")

MF_list = list()
VP_list = list()


for (i in 1:12){
```

```
  m =  models[[i]]

  #Explanatory power
  preds = computePredictedValues(m, expected=FALSE)
  MF = evaluateModelFit(hM=m, predY=preds)

  MF_list[[i]] = MF

  #variance partitioning
  round(head(m$X),2)
  groupnames=c("treatment", "time", "treatment:time") #set variables to groups
  group=c(1,2,2,1,1,3,3,3,3)
  VP = computeVariancePartitioning(m, group=group, groupnames=groupnames)

  VP_list[[i]] = VP
}
```

##### 1.6 Species-specific predictions

```
library(Hmsc)

#all models as a list
load("models_thin_1000_samples_250_chains_4.RData")

#the data where scaling of original data has been done
load("allData_subsets_for_modelling.R")

################################################################################

fms = c(1,3,5,7,9,11)

for(fm in fms){  #loop for each type
  m =  models[[fm]] #occupancy model
  m.abuc = models[[fm+1]] #abundance model

  #matrix for inserting the responses
  responses = data.frame(matrix(0, nrow=length(m[["spNames"]]), ncol=26))
  colnames(responses) <- c("type", "species",
                          "start.rp_median", "start.rp_pp",
                          "start.dp_median","start.dp_pp",
                          "rest.drain_median","rest.drain_pp",
                          "start.rd_median","start.rd_pp",
                          "diff.pris_median", "diff.pris_pp",
                          "abc.drain_median", "abc.drain_pp",
                          "drain.prist.c_median","drain.prist.c_pp")


  for(n in 1:length(m[["spNames"]])){ #loop for each species
    mi=min(m$XData$time)
    ma=max(m$XData$time)
    times = mi:ma
```

```
nt = length(times)
treatments = levels(m$XData$treatment)

for(ft in 1:3){ #loop for each treatment
  XDataNew = data.frame(time = times,
                        treatment = treatments[ft],
                        type =  as.factor(rep(m$XData$type[1],nt)))
  xlev = lapply(m$XData,levels)[unlist(lapply(m$XData,is.factor))]
  XDataNew$type = as.factor(XDataNew$type)
  X = model.matrix(m$XFormula,XDataNew,xlev=xlev)

  post = poolMcmcChains(m$postList)
  post.abuc = poolMcmcChains(m.abuc$postList)
  predN = length(post)
  if(!(predN==length(post.abuc))) print("different length of MCMC chain")
  abu.post = matrix(NA,nrow = predN, ncol=nt)

  for(i in 1:predN){ #loop for each prediction
    L = X%*%post[[i]]$Beta
    p = pnorm(L)
    L = X%*%post.abuc[[i]]$Beta

    if(fm==1) Y.iab <- Y.iab.SL
    if(fm==3) Y.iab <- Y.iab.SH
    if(fm==5) Y.iab <- Y.iab.PL
    if(fm==7) Y.iab <- Y.iab.PH
    if(fm==9) Y.iab <- Y.iab.FL
    if(fm==11) Y.iab <- Y.iab.FH

    #back-transform to original scale
    abuc.tmp <- t(apply(L, 1, function(r)r*attr(Y.iab,'scaled:scale') +
                          attr(Y.iab, 'scaled:center')))

    abuc = exp(abuc.tmp)
    p.sp = p[,n]
    abuc.sp = abuc[,n]
    abu.post[i,] = (p.sp*abuc.sp)
  }

  if(ft==1) start.drained = abu.post[,1]
  if(ft==1) end.drained = abu.post[,nt]
  if(ft==1) change.drained = abu.post[,nt]-abu.post[,1]

  if(ft==2) start.pristine = abu.post[,1]
  if(ft==2) end.pristine = abu.post[,nt]
  if(ft==2) change.pristine = abu.post[,nt]-abu.post[,1]

  if(ft==3) start.restored = abu.post[,1]
  if(ft==3) end.restored = abu.post[,nt]
  if(ft==3) change.restored = abu.post[,nt]-abu.post[,1]
}

responses$type<-paste(m$XData$type[1])
```

```
    responses$species[n] = paste(m[["spNames"]][n])

    #1a Response to drainage (restored-pristine)
    start.rp = start.restored - start.pristine
    responses$start.rp_median[n] <- median(start.rp)
    responses$start.rp_pp[n] <- mean(start.rp>0)

    #1b Response to drainage (drained-pristine)
    start.dp = start.drained - start.pristine
    responses$start.dp_median[n] <- median(start.dp)
    responses$start.dp_pp[n] <- mean(start.dp>0)

    #2 Response to restoration
    rest.drain = change.restored-change.drained
    responses$rest.drain_median[n] = median(rest.drain)
    responses$rest.drain_pp[n] = mean(rest.drain>0)

    #3 Initial difference between restored and drained
    start.rd = start.restored-start.drained
    responses$start.rd_median[n] <- median(start.rd)
    responses$start.rd_pp[n] <- mean(start.rd>0)

    #4 End difference between restored and pristine
    diff.pris = end.restored - end.pristine
    responses$diff.pris_median[n] = median(diff.pris)
    responses$diff.pris_pp[n] = mean(diff.pris>0)

    #5 Species abundance change in drained sites
    abc.drain = change.drained
    responses$abc.drain_median[n] = median(abc.drain)
    responses$abc.drain_pp[n] = mean(abc.drain>0)

    #6 Change in drained sites (in relation to pristine)
    drain.prist.c = abs(end.pristine-end.drained)-abs(start.pristine-start.drained)
    responses$drain.prist.c_median[n] = median(drain.prist.c)
    responses$drain.prist.c_pp[n] = mean(drain.prist.c>0)

  }

  if(fm==1) responses.sl = responses
  if(fm==3) responses.sh = responses
  if(fm==5) responses.pl = responses
  if(fm==7) responses.ph = responses
  if(fm==9) responses.fl = responses
  if(fm==11) responses.fh = responses

}

response.list = list(responses.sl, responses.sh, responses.pl,responses.ph,
                     responses.fl)
```

#### 2. Model-based ordination

##### 2.1 Data

Prior to the model-based ordination, we considered the possible misindentification of the certain Sphagnum species (see Table S1 in Supportin Information 1). For each site separately, we observed which of the species belonging to group have been identified. We assumed that the correct species have been identified at least one of the sampling times. Then, we calculated the total cover of the group for each sample (site_plot_time) and divided this equally between all species belonging to the group which had been observed during the sampling.

```
# Abundance Matrix of species (percent cover)
Y
# Matrix of covariates: plot, site, year, type, treatment, productivity, time
XData
# Data matrix of species information, include species names and
# 'group' to identify moss species and vascular plants
TRData

# General ---------------------------------------------------------------------
# Create numeric time
# Order data according to site,plot and time
Y = Y[with(XData, order(site,plot,timenum)),]
XData = XData[with(XData, order(site,plot,timenum)),]

# Transform percentages to proportions
Ysp = Y/100

# Select species that were observed at least 10 times
Ysp = Ysp[, colSums(Ysp>0)>9]

# relevel treatment
XData$treatment <- relevel(XData$treatment, ref = "pristine")
XData$site = factor(XData$site)
# Create numeric time
XData$timenum = as.numeric(as.character(XData$time))
# Create factor indicating unique combinations of site and monitoring year
XData$siteYear = factor(paste(XData$site,XData$timenum,sep = "_"),
                        levels = unique(paste(XData$site,XData$timenum,
                                              sep = "_")));

# Create a matrix indicating study design
studyDesign = data.frame(plot=factor(XData$plot), site=factor(XData$site),
                         sYear=factor(XData$siteYear))

# There is only few percent covers of 100% and beta distibution only handles
# values (0,1), so ones are replaced with 0.999
Ysp[Ysp == 1] = 0.999
```

##### 2.2 Model fitting

The ordination model is fitted with R package **gllvm**. We fit a generalized linear latent variable model with two latent variables unique to site and monitoring year, species specific intercepts and beta hurdle response model. The shape parameter of beta distribution is set to be common for

```
library(gllvm)
dispGroupall = as.numeric(factor(TRData$group))[TRData$name %in% colnames(Ysp)]
trsXb00_allbh<-system.time(fit_sp_allbh <- try(gllvm(Ysp,
                                                     family = "betaH",
                                                     num.lv = 2,
                                                     studyDesign = studyDesign,
                                                     lvCor = ~(1|sYear),
                                                     starting.val = "zero",
                                                     sd.errors=FALSE,
                                                     optimizer="nlminb",
                                                     disp.formula = dispGroupall )))
```

Gradients are used to check model convergence.

```
gr1 <- c(fit_sp_allbh$TMBfn$gr(fit_sp_allbh$TMBfn$par));
names(gr1) = names(fit_sp_allbh$TMBfn$par)
plot(gr1)
```

Ordination is obtained by plotting latent variables.

```
plot(fit_sp_allbh$lvs)
```

##### 2.3 Predictions

Calculate explanatory power.

```
# Fitted models
fit_sp_allbh

# Pseudo R2's:
# Occupancy 0-1
Y01 = (Ysp>0)*1

# Calculate linear predictors:
LpredLV = (cbind(rep(1, nrow(Ysp)),
                 fit_sp_allbh$X.design,
                 fit_sp_allbh$TMBfn$env$data$dLV%*%fit_sp_allbh$lvs) %*%
             t(cbind(fit_sp_allbh$params$beta0,
                                                  fit_sp_allbh$params$Xcoef,
                     fit_sp_allbh$params$theta%*%diag(fit_sp_allbh$params$sigma.lv,2)) ))

# Tjur's R2 for occupancy
# Prediction probabilities for (Y>0)'s
prob01LV = pnorm(LpredLV[,-(1:178)])
tjurR2 <- function(newy, predi, og=NULL){
  tjurR20<-function(newy, predi){
    (mean(predi[newy==1], na.rm = T) - mean(predi[newy==0], na.rm = T))
  }
  if(is.null(og)) og=1:NCOL(newy)
  tjurog<-NULL
```

```
  for (i in unique(og)) {
    tjurog[i]=tjurR20(newy[, og==i],predi[, og==i])
  }
  tjurog
}
# species specific Tjur's R2 measures
tjrsx = tjurR2(Y01, prob01LV)

# Linear predictors for Cover model
LinPredLV = LpredLV[,1:178]

R2lvi=NULL
og = (1:ncol(Ysp_allf))
for (i in 1:max(og)) {
  newyk = (Ysp_allf[, og==i])[Y01[,og==i]>0]
  predik = (LinPredLV[, og==i])[Y01[,og==i]>0]
  if(length(unique(newyk))>2 | length(unique(predik))>2) try(R2lvi[i] <-
                                                             cor(qnorm(newyk), predik)^2)
}
FerrariCribariNetoR2lvi = R2lvi

# Plot
pseudoR2meas = cbind(tjrsx, FerrariCribariNetoR2lvi)
colnames(pseudoR2meas) = c("occupancy", "cover")

library(ggplot2)
datalong1 =data.frame(pseudoR2 = c(pseudoR2meas),
                      Ptype= factor(rep(colnames(pseudoR2meas),
                                        each = nrow(pseudoR2meas))))

plot0 <- ggplot(data = datalong1, aes(x=factor(Ptype), y=pseudoR2)) +
  geom_boxplot(fill="grey") +
  xlab("") +
  ylab(expression("Pseudo R"^2)) + ylim(0,1)+
  theme_bw()
plot0
```

Predictive powers are evaluated based on 4-fold cross validation.

```
# 4 fold cross validation
# Create folds:
set.seed(111)
sampleidrop = vector(length = nrow(studyDesign))
for (i in levels(studyDesign$sYear)) {
  si = studyDesign$sYear == i
  if((sum(si)-4)>0){
    s4_ = sample(1:4,sum(si)-4, replace = TRUE)
    sampleidrop[si] = sample(c(1:4,s4_),size= sum(si))
  } else {
    sampleidrop[si] = 5
  }
}
table(sampleidrop)
```

```r
sampleidrop4=sampleidrop
sampleidrop4[sampleidrop4==5]=4

# Calculate predictions
LinPredM = matrix(NA, nrow(Ysp), ncol(Ysp)*2)
for (i in 1:4) {
trsXb00_allbh2_drop1<-system.time(fiti <- try(gllvm(Ysp[sampleidrop!=i,],
                                                    family = "betaH",
                                                    num.lv = 2,
                                                    studyDesign = studyDesign[sampleidrop!=1,],

                                                    lvCor = ~(1|sYear),
                                                    Lambda.struc="diagonal",
                                                    sd.errors=FALSE,
                                                    disp.formula = dispGroupall )))
  Ypred = Ysp[sampleidrop4==i,]
  Ypred01=(Ypred>0)*1
  SDe = studyDesign[sampleidrop4==i,]
  Dsyear =  model.matrix(~SDe$sYear-1)
  LinPredi = Dsyear%*%(cbind(1,fiti$lvs)%*%rbind(fiti$params$beta0,
                                                t(fiti$params$theta %*%
                                                 diag(fiti$params$sigma.lv,2))))
  LinPredM[sampleidrop4==i,] = LinPredi
}

# Calculate pseudo R2's
# Occupancy
Y01pred = (Ysp>0)*1
prob01pred = pnorm(LinPredM[,-(1:178)])
predTR2 = tjurR2(Y01pred, prob01pred)

# Cover
LinPred = LinPredM[,1:178]

R2lviPred=NULL
og = (1:ncol(Ysp))
for (i in 1:max(og)) {
  newyk = (Ysp[, og==i])[Y01pred[,og==i]>0]
  predik = (LinPred[, og==i])[Y01pred[,og==i]>0]
  # R2 by Ferrari and Cribari-Neto (2004, p. 806)
  if(length(unique(newyk))>2 | length(unique(predik))>2) try(R2lviPred[i] <-
                                                            cor(qnorm(newyk),
                                                               predik)^2)
}
```
